# Multivalent lipid targeting by the calcium-independent C2A domain of Slp-4/granuphilin

**DOI:** 10.1101/2020.05.29.123596

**Authors:** Aml A Alnaas, Abena Watson-Siriboe, Sherleen Tran, Mikias Negussie, Jack A. Henderson, J. Ryan Osterberg, Nara Lee Chon, Julianna Oviedo, Tatyana Lyakhova, Cole Michel, Nichole Reisdorph, Richard Reisdorph, Colin T. Shearn, Hai Lin, Jefferson D. Knight

## Abstract

Synaptotagmin-like protein 4 (Slp-4), also known as granuphilin, is a Rab effector responsible for docking secretory vesicles to the plasma membrane before exocytosis. Slp-4 binds vesicular Rab proteins via an N-terminal Slp homology (SHD) domain, interacts with plasma membrane SNARE complex proteins via a central linker region, and contains tandem C-terminal C2 domains (C2A and C2B) with affinity for phosphatidylinositol-(4,5)-bisphosphate (PIP_2_). The Slp-4 C2A domain binds with low nanomolar apparent affinity to PIP_2_ in lipid vesicles that also contain background anionic lipids such as phosphatidylserine (PS), but much weaker when either the background anionic lipids or PIP_2_ are removed. Through computational and experimental approaches, we show that this high affinity membrane interaction arises from concerted interaction at multiple sites on the C2A domain. In addition to a conserved PIP_2_-selective lysine cluster, there exists a larger cationic surface surrounding the cluster which contributes substantially to the affinity for physiologically relevant lipid compositions. While mutations at the PIP_2_-selective site decrease affinity for PIP_2_, multiple mutations are needed to decrease binding to physiologically relevant lipid compositions. Docking and molecular dynamics simulations indicate several conformationally flexible loops that contribute to the nonspecific cationic surface. We also identify and characterize a covalently modified variant in the bacterially expressed protein, which arises through reactivity of the PIP_2_-binding lysine cluster with endogenous bacterial compounds and has a low membrane affinity. Overall, multivalent lipid binding by the Slp-4 C2A domain provides selective recognition and high affinity docking of large dense-core secretory vesicles to the plasma membrane.

---

Synaptotagmin like protein 4 (Slp-4), also known as granuphilin, plays an important role in the trafficking and docking of insulin and other large dense core secretory vesicles to the plasma membrane prior to exocytosis (1). Two seemingly contradictory effects have been observed upon overexpression of Slp-4 in insulin-secreting Min6 and INS-1 cell lines: (i) an increased number density of secretory granules docked to the plasma membrane prior to stimulation (2,3), but (ii) decreased efficiency of secretion, and a decreased total amount of insulin secreted upon stimulation with KCl or glucose (3,4). Inversely, knockout or knock-down of Slp-4 enhances insulin secretion (5,6). The precise intermolecular interactions which give rise to these effects are not yet clear.

Slp-4 is classified as a Rab effector protein which targets to secretory vesicles via interaction of its N-terminal Slp homology domain (SHD) with vesicular Rab27a, for which it recognizes both GDP and GTP-bound forms (7). The central linker region of Slp-4 interacts with plasma membrane SNARE complex proteins (8-11). The protein’s tandem C-terminal C2 domains (C2A and C2B) bind anionic plasma membrane lipids including phosphatidylinositol-(4,5)-bisphosphate (PIP_2_) in a Ca^2+^-independent manner (12,13). The interactions between Slp family members and Rab proteins are well studied; for example, point mutations in the Slp-4 SHD domain that prevent binding to Rab27 are known (3,7,12,14). However, the interactions that allow Slp-4 to bridge to the plasma membrane, including those of the C2 domains, are less thoroughly characterized.

The Slp-4 C2A domain plays a critical role in targeting its parent protein to the plasma membrane. Overexpression of a ΔC2AB mutant leads to fewer vesicles docked to the plasma membrane as compared to the full length Slp-4, and the mutant protein localizes to the cytosol and interior vesicles rather than to docked vesicles (6,12). However, deletion of only the C2B domain has little effect; indeed, the Slp-4b splice variant lacks a C2B domain but has similar functionality to the full-length Slp-4a version (4,6,10).

Multivalent protein-membrane interactions are abundant among membrane-targeting proteins, and have been suggested to constitute the principal mechanism for achieving specificity in targeting to particular subcellular membranes (15,16). Our group has previously shown that the Slp-4 C2A domain binds with apparent low nanomolar affinity to PIP_2_ in physiological liposome compositions, while the affinity of the C2B domain towards the same liposomes is approximately 100-fold weaker (13). Therefore, the C2A domain is likely the most important mediator of Slp-4 plasma membrane interaction. Interestingly, the Slp-4 C2A domain binds with only modest (∼1 µM) affinity to PIP_2_ in liposomes with a phosphatidylcholine (PC) background, or to soluble PIP_2_ (17). This discrepancy suggests that besides the PIP_2_ binding site, there must be other lipid contacts that stabilize the association of the Slp-4 C2A domain to lipid compositions resembling the plasma membrane.

The available crystal structure of the Slp-4 C2A domain (PDB: 3FDW) shows an 8-stranded β-sandwich structure that aligns well with other C2 domains with type I topology, including those from synaptotagmins and other related proteins (18) (**Figure 1**). It is missing four of the five conserved Asp residues in the β2-β3 and β6-β7 loops that coordinate Ca^2+^ in several well-studied C2 domain proteins including synaptotagmins (18,19). However, it contains a PIP_2_-binding sequence including a cluster of lysine residues on the β3 and β4 strands, also termed the polybasic region, which is found in other PIP_2_-binding type I topology C2 domains including protein kinase Cα, the C2B domain of synaptotagmin-1, and the C2A domains of rabphilins (16). The PIP_2_ binding site of Slp-4 C2A was previously reported to center on this cluster (17). Notably, a large region around this cluster contains other basic residues that impart a positively charged electrostatic surface (**Figure 1**).

**Figure 1.**
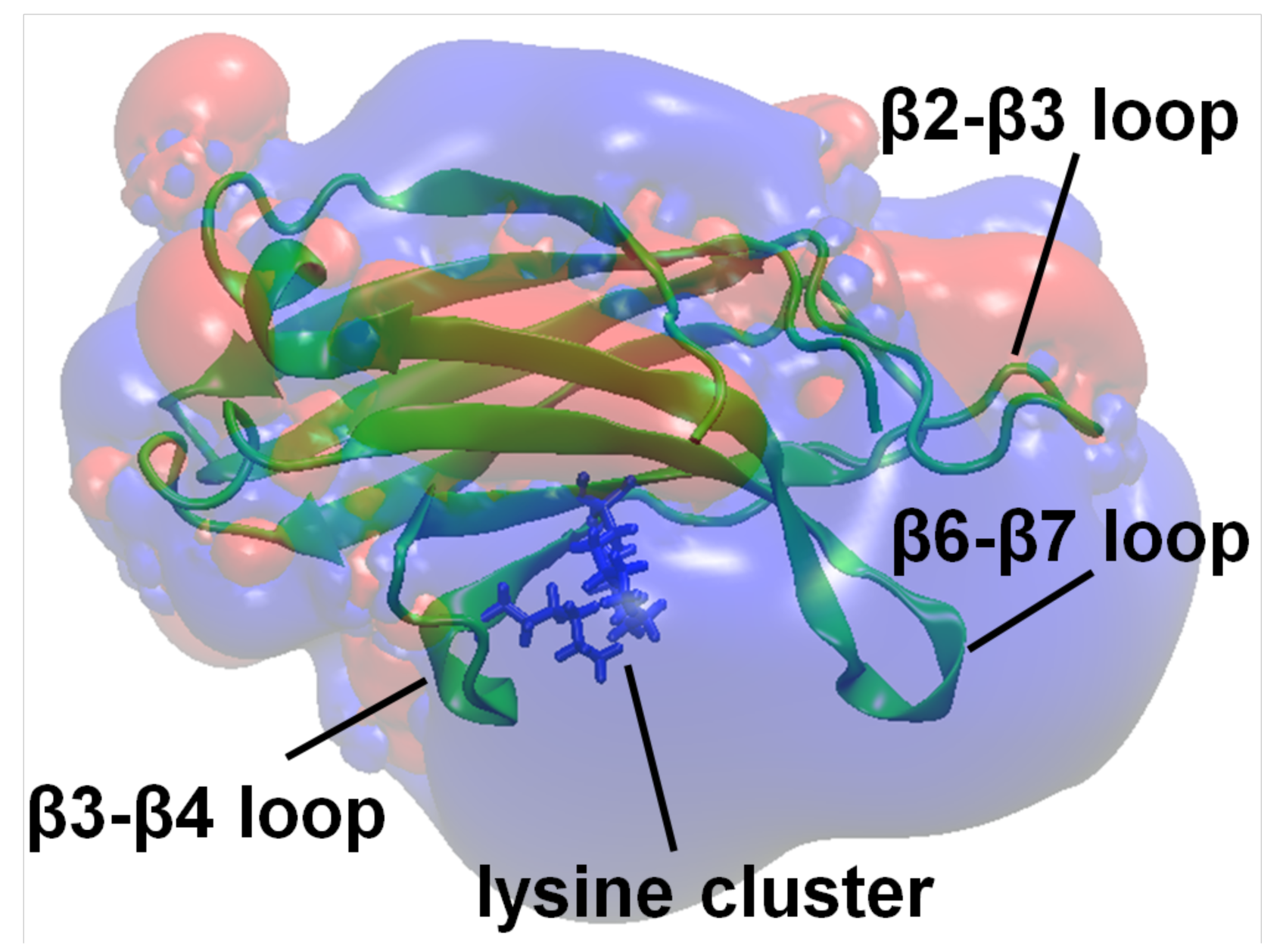
Slp-4 C2A domain structure and electrostatic surface. The locations of important loops and the PIP_2_-binding lysine cluster are shown, along with an electrostatic surface map showing the large positive (blue) surface surrounding the lysine cluster.

In this work we have sought to identify how background anionic plasma membrane lipids, such as phosphatidylserine (PS) and phosphatidylinositol (PI), work in conjunction with PIP_2_ to mediate strong membrane binding of the Slp-4 C2A domain. Using purified C2A domains and synthetic liposomes with controlled lipid compositions, we have tested the contributions of PIP_2_ and other anionic lipids to liposome affinity as well as kinetic on- and off-rates. As a complement to these experiments, we have used computational docking and molecular dynamics simulations to predict which regions of the polycationic protein surface interact with each target lipid headgroup or insert into the hydrophobic interior of a lipid bilayer. Finally, we have tested the effect of single and multiple point mutations on the C2A domain’s membrane binding using both experimental and computational biochemistry. Overall, we find that binding to PIP_2_ occurs primarily through the previously identified lysine cluster (17), while interaction with anionic background lipids involves a large surface surrounding this lysine cluster and encompassing multiple loop regions. These interactions significantly enhance the membrane affinity, and their broad surface distribution renders membrane binding relatively insensitive to effects of mutations. Furthermore, we show for the first time that the conserved lysine cluster is susceptible to covalent modification by carbonyl-containing compounds during bacterial protein expression, and we characterize the effect of this modification on protein-membrane binding.

## Results

### Strong Slp-4 C2A membrane binding requires PS and PIP_2_

The C2A domain of Slp-4 is known to drive this protein’s ability to bind cellular plasma membranes primarily via interaction with PIP_2_ (12,13). However, the protein retains an affinity for membranes containing the anionic lipids PS and/or PI, even in the absence of PIP_2_ (13). Here, we set out to discern the contributions of PIP_2_ and background anionic lipid binding to its strong affinity for physiological lipid membranes, including determining residues that drive binding to anionic background lipids. In order to do this, we first measured interactions between purified protein domains and liposomes of defined lipid composition (**Table 1**).

**Table 1.**
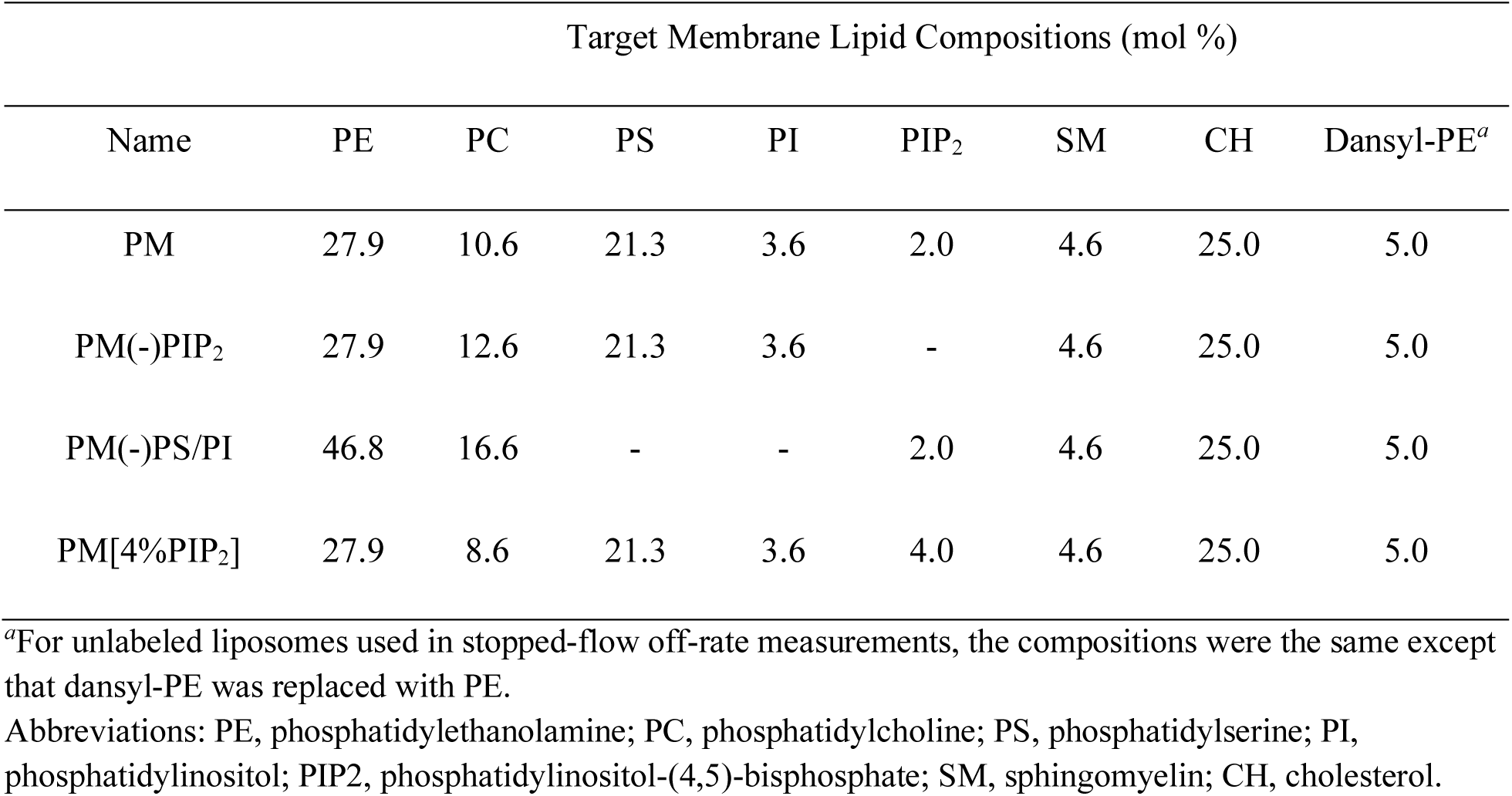
Lipid compositions used in this study.

| Target Membrane Lipid Compositions (mol %) |  |  |  |  |  |  |  |  |
| --- | --- | --- | --- | --- | --- | --- | --- | --- |
| Name | PE | PC | PS | PI | PIP<sub>2</sub> | SM | CH | Dansyl-PE<sup>a</sup> |
| PM | 27.9 | 10.6 | 21.3 | 3.6 | 2.0 | 4.6 | 25.0 | 5.0 |
| PM(-)PIP<sub>2</sub> | 27.9 | 12.6 | 21.3 | 3.6 | - | 4.6 | 25.0 | 5.0 |
| PM(-)PS/PI | 46.8 | 16.6 | - | - | 2.0 | 4.6 | 25.0 | 5.0 |
| PM[4%PIP<sub>2</sub>] | 27.9 | 8.6 | 21.3 | 3.6 | 4.0 | 4.6 | 25.0 | 5.0 |

<sup>a</sup>For unlabeled liposomes used in stopped-flow off-rate measurements, the compositions were the same except that dansyl-PE was replaced with PE.
Abbreviations: PE, phosphatidylethanolamine; PC, phosphatidylcholine; PS, phosphatidylserine; PI, phosphatidylinositol; PIP<sub>2</sub>, phosphatidylinositol-(4,5)-bisphosphate; SM, sphingomyelin; CH, cholesterol.

While the Slp-4 C2A domain has a strong affinity for liposomes with a lipid composition approximating the plasma membrane interior leaflet, removal of either PIP_2_ or background anionic lipids (PS and PI) decreases its affinity by an order of magnitude (**Figure 2A**). We measured the relative affinities of the C2A domain for various synthetic liposomes containing dansyl-PE lipids by measuring tryptophan-to-dansyl FRET of the lipid-bound Slp-4 C2A domain while titrating with soluble inositol-(1,2,3,4,5,6)-hexakisphosphate (IP_6_), which we have previously demonstrated inhibits lipid binding by this protein (13). For a given concentration of liposomes and protein, the concentration of inhibitor required to remove 50% of the protein from the membrane (IC_50_) is approximately proportional to the mole-fraction equilibrium constant *K*_x_ for partitioning onto the liposome surface (Eq. 2, see Methods). When liposomes were used approximating the plasma membrane inner leaflet lipid composition (PM, **Table 1**), the IC_50_ of IP_6_ was 620 ± 80 µM, corresponding to a binding constant of (30 ± 5) × 10^7^ (**Figure 2A, Table 2**). Consistent with our previous report (13), removal of PIP_2_ from the liposome lipid composition [PM(-)PIP_2_] decreased the IC_50_ and the membrane partitioning coefficient by a factor of ∼30 (**Table 2**). Here we also show that removal of background anionic lipids PS and PI while maintaining 2% PIP_2_ in the liposomes [PM(-)PS/PI] also significantly decreased the IC_50_, reflecting a ∼12-fold decrease in the affinity (**Figure 2A**; **Table 2**). The membrane binding is primarily electrostatic, as binding to all three liposome compositions could be screened by NaCl at 2-3x physiological ionic strength (**Figure 2B**). Increased salt concentration more efficiently screened binding to liposomes lacking PIP_2_ or background anionic lipids, relative to liposomes containing both. Thus, physiologically relevant levels of both PIP_2_ and background anionic lipids make important contributions to membrane binding for this protein.

**Table 2.**
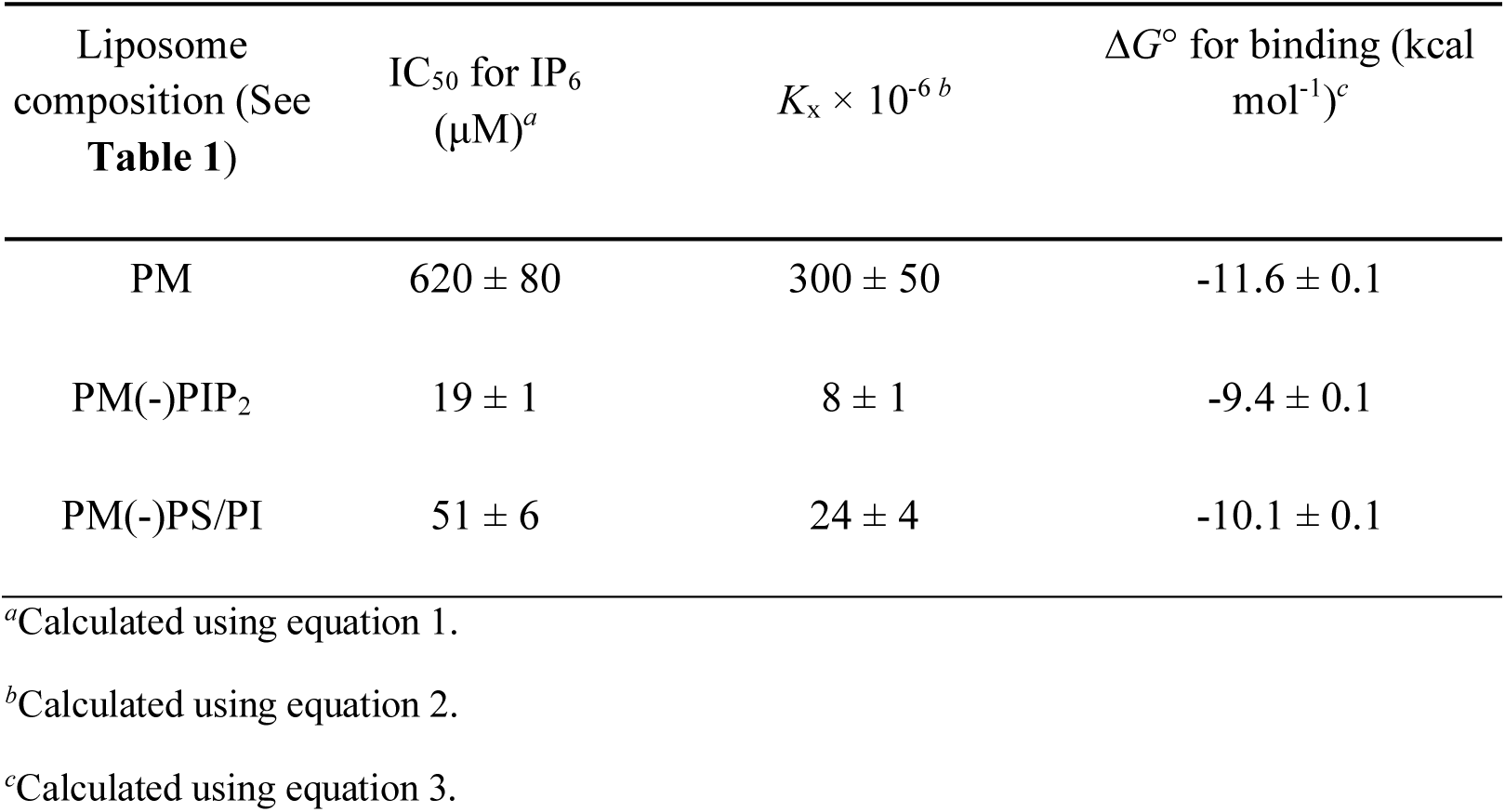
Equilibrium IP_6_ titration parameters of wild-type Slp-4 C2A domain

| Liposome composition (See **Table 1**) | IC<sub>50</sub> for IP<sub>6</sub> (μM)<sup>a</sup> | $K_x \times 10^{-6}$<sup>b</sup> | ΔG° for binding (kcal mol<sup>-1</sup>)<sup>c</sup> |
| --- | --- | --- | --- |
| PM | 620 ± 80 | 300 ± 50 | -11.6 ± 0.1 |
| PM(-)PIP<sub>2</sub> | 19 ± 1 | 8 ± 1 | -9.4 ± 0.1 |
| PM(-)PS/PI | 51 ± 6 | 24 ± 4 | -10.1 ± 0.1 |

<sup>a</sup>Calculated using equation 1.
<sup>b</sup>Calculated using equation 2.
<sup>c</sup>Calculated using equation 3.

**Figure 2.**
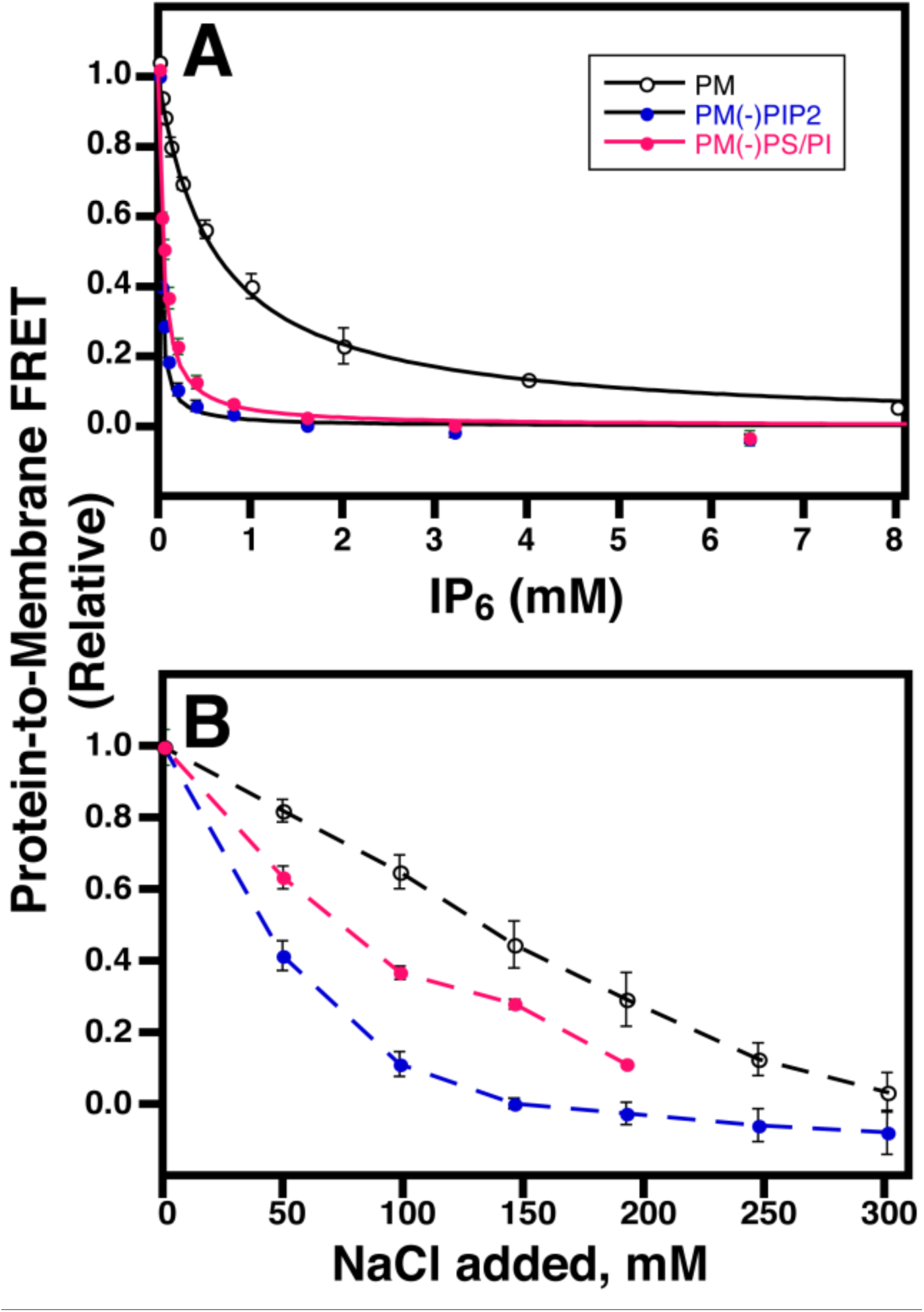
Lipid dependence of Slp-4 C2A domain membrane binding and anion inhibition. **A:** IP_6_ competition titrations for Slp-4 C2A initially bound to liposomes of the indicated compositions. Best-fit curves to a competitive inhibition binding model (Eq. 2) are shown; IC_50_ values are given in **Table 2. B:** NaCl screening titrations. Dashed lines guide the eye. Error bars are standard deviation of 3 independent replicate titrations and where not visible are smaller than the data points. Colors represent different liposome compositions (**Table 1**): black, PM; blue, PM(-)PIP_2_; red, PM(-)PS/PI.

Stopped-flow kinetic measurements of Slp-4 C2A domain liposome dissociation also show a pronounced dependence on both PIP_2_ and background anionic lipids. We previously reported that removal of PIP_2_ increases the membrane dissociation rate of the domain while having no significant effect on the association rate (13). Here, we show that removal of background anionic lipids also significantly increased the kinetics of dissociation (**Figure 3; Table 3**, *k*_off_) while having little effect on the association rate constant (**Table 3**, *k*_on,x_). The strong dependence of dissociation rates on both PIP_2_ and background anionic lipids indicates that both species contribute significantly to the thermodynamic stability of the membrane-bound state. Furthermore, the affinity differences among the various lipid compositions tested are due almost entirely to the differences in off rate. Dissociation from all liposome compositions was revealed to be biexponential, suggesting the presence of multiple membrane-bound states, which could reflect populations associated with different numbers of bound lipids. This feature was not apparent in our previous report due to a smaller signal-to-noise ratio (13).

**Table 3.** Stopped-flow kinetic parameters of wild-type Slp-4 C2A domain

| Liposome composition<br>(See Table 1) | $k_{\text{obs}}$ ($\text{s}^{-1}$) | $k_{\text{on,x}}$ ($\text{s}^{-1}$) $\times 10^{-6a}$ | $k_{\text{off}}$ ($\text{s}^{-1}$) |
| --- | --- | --- | --- |
| PM | $30 \pm 12$ | $43 \pm 18$ | $1.0 \pm 0.3$ ($35 \pm 18\%$ amp) |
| | | | $0.14 \pm 0.03$ ($65 \pm 18\%$ amp) |
| PM(-)PIP<sub>2</sub> | $90 \pm 50$ | $90 \pm 60$ | $21 \pm 13$ ($66 \pm 13\%$ amp) |
| | | | $2.6 \pm 1.3$ ($34 \pm 13\%$ amp) |
| PM(-)PS/PI | $55 \pm 15$ | $73 \pm 23$ | $8.0 \pm 1.5$ ($60 \pm 15\%$ amp) |
| | | | $1.3 \pm 0.4$ ($40 \pm 15\%$ amp) |
| PM/4%PIP<sub>2</sub> | $47 \pm 3$ | $69 \pm 8$ | $0.43 \pm 0.14$ ($27 \pm 19\%$ amp) |
| | | | $0.08 \pm 0.05$ ($73 \pm 19\%$ amp) |

<sup>a</sup>Calculated using equation 7.

**Figure 3.**
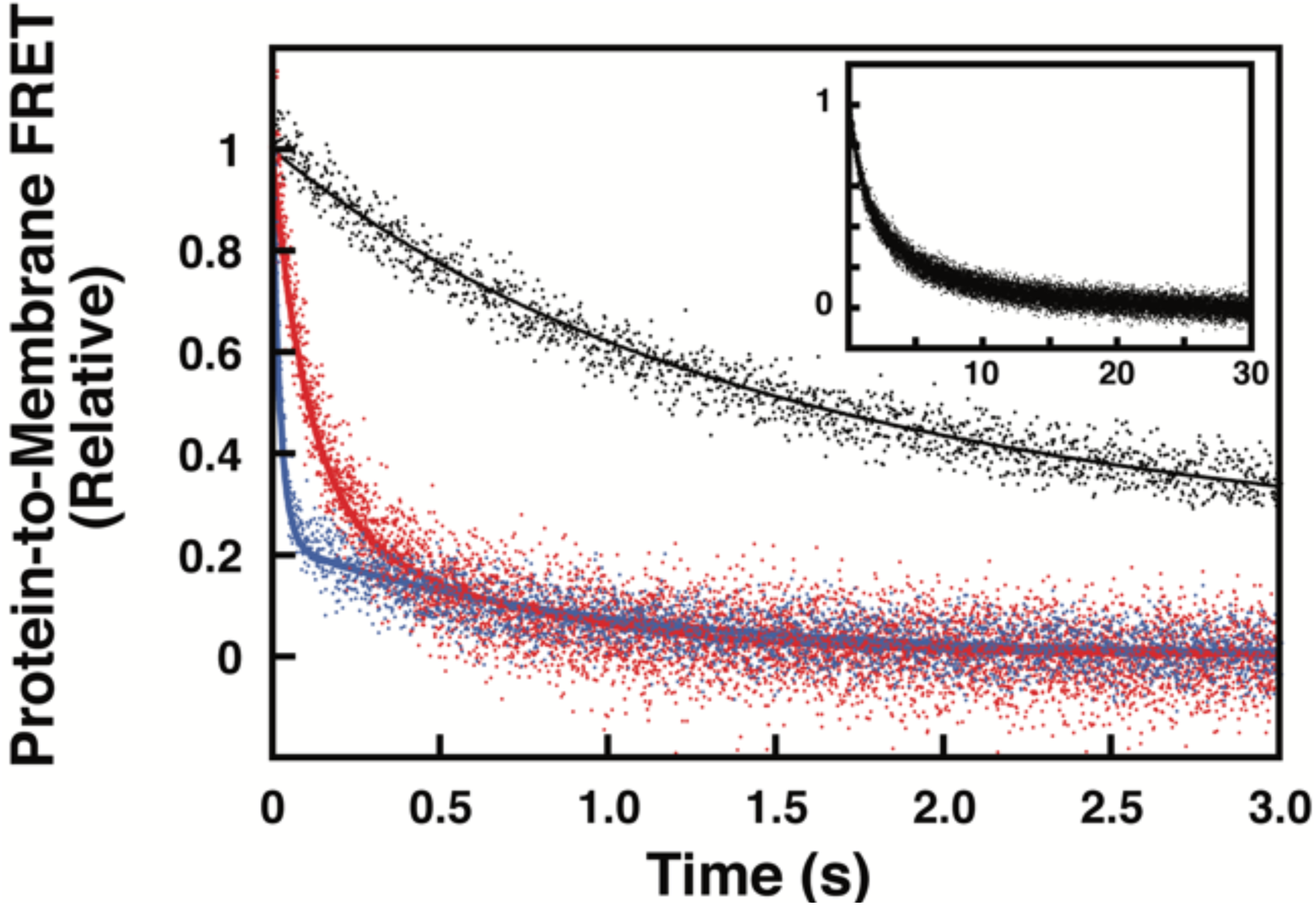
Lipid composition dependence of Slp-4 C2A domain dissociation kinetics. Representative dissociation curves are shown from PM (black), PM(-)PIP_2_ (blue), and PM(-)PS/PI (red) liposomes (See **Table 1** for lipid compositions). **Inset:** full time course of PM dissociation curve.

Increasing the PIP_2_ concentration in the PM liposome composition from 2% to 4% further slowed dissociation, by about a factor of 2 relative to PM (**Table 3**). This factor is much smaller than the ∼10-fold difference in off-rate comparing PM(-)PIP_2_ to PM compositions, suggesting that the increased PIP_2_ concentration allows engagement of weaker, secondary binding site(s) on the protein. This secondary binding appears to be nonselective, as phosphatidic acid (PA) had a similar effect: dissociation from liposomes containing 2% PIP_2_ and 2% PA proceeded on a timescale comparable to PM/4%PIP_2_ (**Figure S1; Table S1**). In contrast, the primary PIP_2_ binding is highly selective for PIP_2_ over PA, as dissociation from liposomes containing 2% PA but not PIP_2_ was much faster than from PM liposomes [**Figure S1 and Table S1**, compare PM(-)PIP_2_(+)PA to PM]. Overall, the results of kinetic experiments confirm that this C2 domain’s strong membrane affinity relies on the presence of both PIP_2_ and background anionic lipids.

### Predicting binding sites for anionic ligands using computational modeling

In order to predict which regions of the Slp-4 C2A domain bind PIP_2_ and or other anionic ligands, we simulated inositol-(1,4,5)-trisphosphate (IP_3_), a soluble PIP_2_ analogue, docking to the protein domain using a molecular docking algorithm. The first simulation included 510 separate docking calculations for IP_3_ molecules placed at a library of positions around the published protein structure (PDB: 3FDW). As expected, the majority of these calculations showed docking to the known PIP_2_ binding site, which is centered on the cluster of conserved lysine residues (K398, K410, and K412) in the β3 and β4 strands (**Figure 4, Lys cluster**) (16,17).

**Figure 4:**
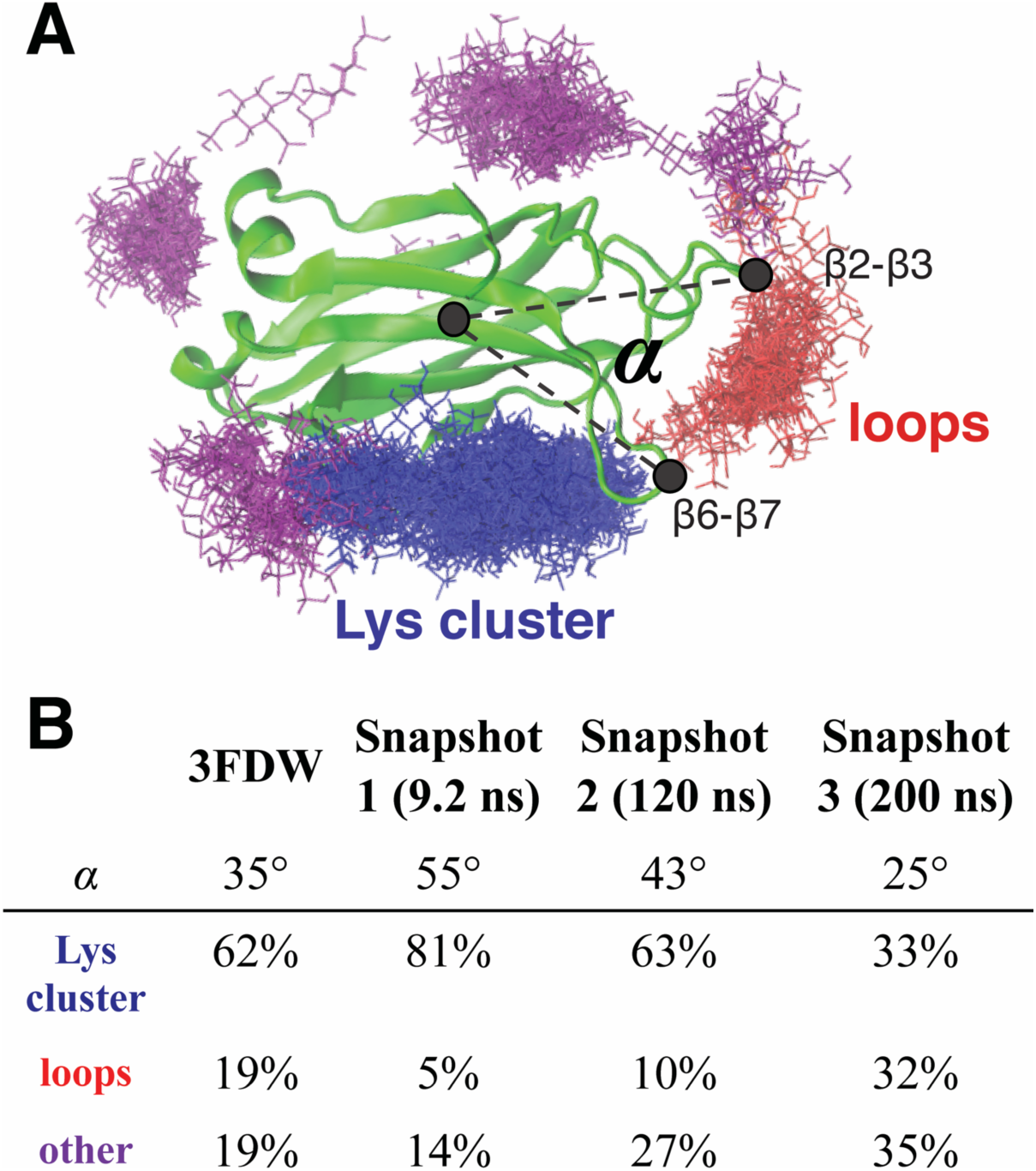
Docking calculations with Slp-4 C2A domain. **A:** Representative clustering of docked IP_3_ ligands following Flexidock calculations. A total of 510 starting configurations were modeled, and all of the final ligand positions are shown overlaid in stick format, colored by location cluster. Gray dashed lines define the angle *α* between the β2-β3 and β6-β7 loops. **B:** Tabulation of *α* angle and ligand clustering for four similar sets of Flexidock calculations, each using a different starting protein conformation from the crystal structure (3FDW) or from the indicated timepoint of the standalone molecular dynamics simulation (**Movie 1**).

Several other clusters emerged from these calculations as secondary docking sites for IP_3_, suggesting they could represent nonspecific binding sites for anionic ligands. The largest of these clusters was near the tips of the β2-β3 and β6-β7 loops, which are structurally homologous to calcium-binding loops 1 and 3 in Ca^2+^-sensitive C2 domains (**Figure 4, loops**) (18). These loop regions are known to be dynamic in Ca^2+^-free C2 domains (20,21); therefore, we conducted a 200-ns molecular dynamics simulation of the protein domain alone in physiological salt solution at pH 7 (Supporting Information, **Movie S1**). As expected, the loop regions showed conformational fluctuations during this simulation while the core of the domain remained intact; in particular, the β6-β7 loop moved closer to the β2-β3 loop during the simulation (**Figure S2**); i.e., the angle α shown in **Figure S2A** and **Figure 4A** decreased over time following an initial rapid increase.

Three snapshots from this simulation with varying α angles were chosen as templates for additional IP_3_ docking calculations. Results of these calculations are tabulated in **Figure 4B** alongside those of the original calculation based on the crystal structure. Docking to the loops site correlated strongly with α: as α decreased during the course of the MD simulation, the percentage of docking events to the primary PIP_2_ binding site decreased while docking to the tips of the loops increased. This result suggests that proximity between these two loops, both of which contain basic residues, improves affinity towards anionic ligands at this site. Further docking calculations using protein variants with mutations in the primary PIP_2_ binding site shifted IP_3_ binding further toward the loops site, consistent with the hypothesis that this is a preferred secondary binding site for anionic ligands (**Table S2**).

### Mapping protein-lipid contacts from MD simulations

Molecular dynamics simulations of the Slp-4 C2A domain docking to anionic membranes indicate that multiple regions of the protein domain, including the β2-β3 and β6-β7 loops, make electrostatic and/or hydrophobic contacts with the membrane. In order to illuminate what residues might make key contacts with target lipids, we performed three parallel molecular dynamics simulations of the Slp-4 C2A domain binding to lipid bilayers composed of PC, PS, and PIP_2_ (**Figure S3; Movies S2-S4**). The three simulations differed in the initial placement of the protein with respect to the two PIP_2_ molecules embedded in the target membrane leaflet. In Models 1 and 2, the protein was placed midway between the two PIP_2_ molecules with its long axis oriented perpendicular to (Model 1) or along (Model 2) the line between the PIP_2_ molecules, while in Model 3 the protein was placed with its lysine cluster directly above one of the PIP_2_ molecules. In all three simulations, the protein rapidly made contact with the lipids immediately below it within the first 20 ns, after which the center-of-mass position of the protein remained relatively constant relative to the membrane phosphate plane (**Figure 5A**). Because of this rapid membrane association and the initial configurations of each system, the lysine cluster only interacted with PIP_2_ in Model 3; in Models 1 and 2, the PIP_2_ molecules interacted with residues at the periphery of membrane contact. This result suggests that membrane rearrangements may occur to allow PIP_2_ insertion into its primary binding site, but likely on a slower timescale than in these simulations. A control simulation was also performed with the protein initially positioned above a PC bilayer (**Figure S3; Movie S5**). In this simulation, the protein initially drifted away from the membrane before making contact with the polar headgroup region via the β3-β4 loop (**Figure 5A**).

**Figure 5:**
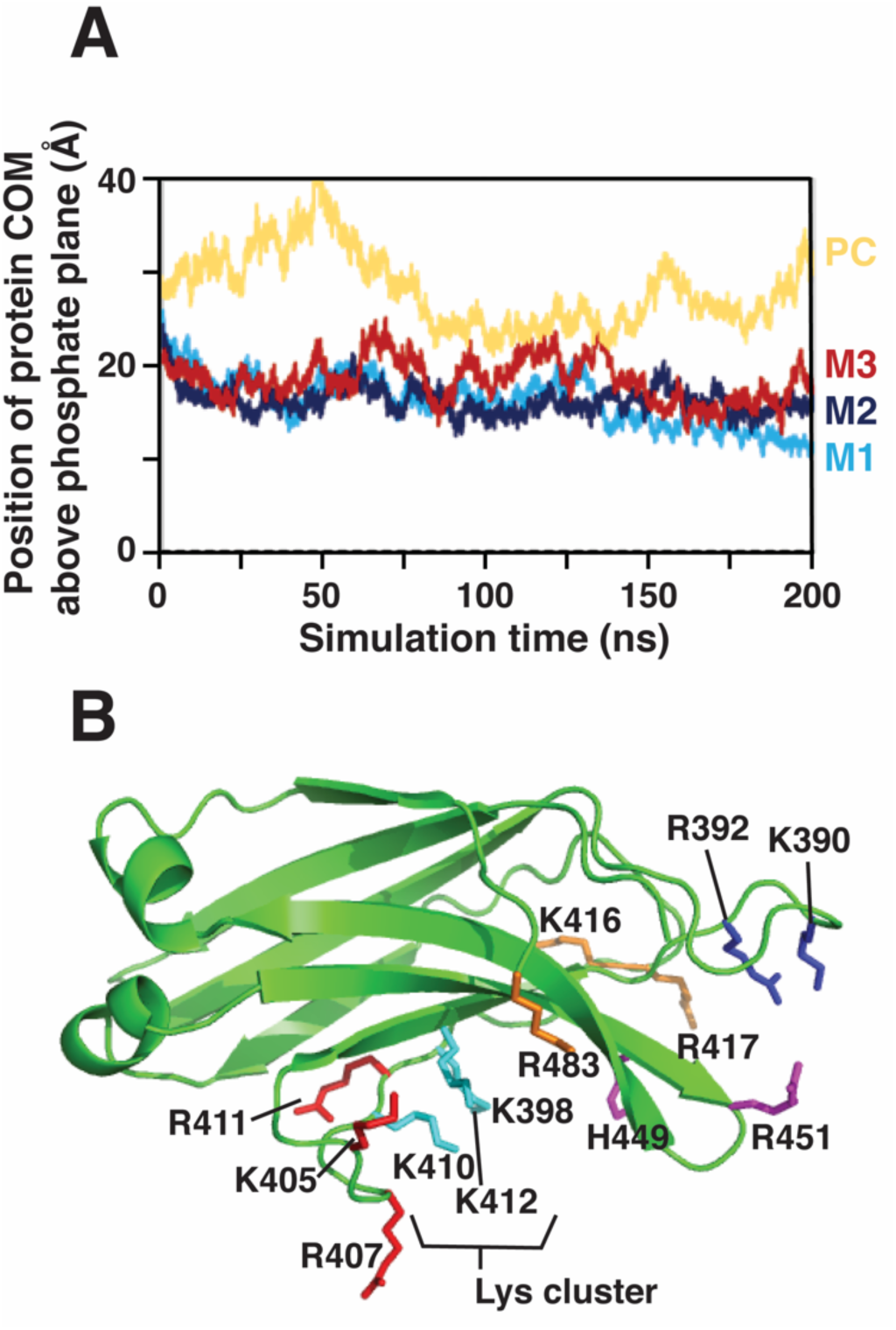
Mapping protein-membrane contacts from MD simulations. **A:** Height of the protein center-of-mass (COM) above the phosphate plane of the membrane during each simulation. Yellow: PC control; light blue: Model 1 (M1); dark blue: Model 2 (M2); red: Model 3 (M3). **B:** Residues that make significant ionic contact with PS or PIP_2_ in at least one of the simulations are labeled. Cyan: conserved lysine cluster; blue: β2-β3 loop; red: β3-β4 loop and β4 strand; purple: β6-β7 loop; orange: β4-β5 loop and C-terminus.

For the PC/PS/PIP_2_ simulations, protein-lipid contacts were identified throughout the polycationic surface of the protein (**Figure 1**). Residues making contact with PS and PIP_2_ included the two regions predicted from the IP_3_ docking calculations: the lysine cluster and the loops region (which includes the β2-β3 loop and the β6-β7 loop) (**Figure 5B; Table 4**). Additionally, basic residues in the β3-β4 loop near the lysine cluster made significant contact. The β2-β3 and β6-β7 loops remained much closer together (small α angle) throughout all of the simulations than in the crystal structure (**Figure S2B-C**). Residues that average ≥ 0.5 contacts with PS or PIP_2_ during the last 100 ns of at least one simulation are quantified in **Table 4**. These include Lys390 and Arg392 in the β2-β3 loop, several residues in the β3-β4 region, and His449 and Arg451 in the β6-β7 loop. Contacts for other basic residues from the MD simulations are listed in **Table S3**. Notably, Lys398 contributed centrally to PIP_2_ coordination in Model 3 (in which the PIP_2_ was bound near the previously reported binding pocket), but did not coordinate PS or PIP_2_ in the other simulations. The lipid contacts of each residue are plotted as a function of simulation time in **Figures S4 and S5**.

**Table 4.** Electrostatic hydrophilic contacts. The number of the indicated lipid molecules (± S.D.) contacted by each residue, averaged over the last 100 ns of simulations, is displayed for the PC control and PC/PS/PIP_2_ Models 1, 2, and 3. Residues listed here had contact numbers ≥ 0.5 in any of the simulations; an extended list including all basic residues is given in **Table S3**. Contacts are defined as described in Methods.

|  |  | PC control | Model 1 |  | Model 2 |  | Model 3 |  |
| --- | --- | --- | --- | --- | --- | --- | --- | --- |
|  |  | Avg PC Contact | Avg PS Contact | Avg PIP<sub>2</sub> Contact | Avg PS Contact | Avg PIP<sub>2</sub> Contact | Avg PS Contact | Avg PIP<sub>2</sub> Contact |
| β2-β3 loop | **K390** | 0 | 1.3 ± 0.8 | 0 | 0.3 ± 0.5 | 0.1 ± 0.0 | 0 | 0 |
| | **R392** | 0 | 1.1 ± 0.0 | 0 | 0.1 ± 0.3 | 1.0 ± 0.0 | 0.5 ± 0.7 | 0 |
| β3-β4 loop | **K405** | 1.8 ± 0.8 | 0 | 0 | 0 | 1.0 ± 0.1 | 0.3 ± 0.5 | 0 |
| | **R407** | 3.0 ± 0.7 | 0.1 ± 0.3 | 0 | 0.2 ± 0.4 | 1.0 ± 0.0 | 0.9 ± 0.5 | 0 |
| Lys cluster | **K398** | 0.3 ± 0.6 | 0 | 0 | 0 | 0 | 0.6 ± 0.5 | 1.0 ± 0.1 |
| | **K410** | 0.1 ± 0.3 | 0 | 0 | 0 | 0 | 0.4 ± 0.5 | 0 |
| β4 | **R411** | 0.4 ± 0.7 | 1.6 ± 0.6 | 0 | 0.2 ± 0.4 | 0 | 0.7 ± 0.5 | 0 |
| Lys cluster | **K412** | 0.2 ± 0.4 | 0.8 ± 0.5 | 0 | 0.1 ± 0.3 | 0 | 0 | 1.0 ± 0.0 |
| β4-β5 loop | **K416** | 0 | 0.3 ± 0.5 | 0 | 0.5 ± 0.5 | 0 | 0 | 0 |
| | **R417** | 0.1 ± 0.4 | 0.2 ± 0.5 | 0 | 1.0 ± 0.6 | 0.1 ± 0.3 | 0.3 ± 0.5 | 0 |
| β6-β7 loop | **H449** | 0.1 ± 0.3 | 0 | 0 | 0.2 ± 0.4 | 0 | 0 | 0.9 ± 0.3 |
| | **R451** | 0 | 0 | 0 | 0.5 ± 0.5 | 0 | 0.5 ± 0.8 | 0 |
| C-term | **K483** | 0.2 ± 0.4 | 0 | 1.0 ± 0.2 | 0 | 0 | 0 | 0 |

Although the membrane contacts of this protein domain are predominantly ionic, there were also uncharged residues on the β3-β4 loop that made extensive contact with the membrane in the MD simulations, including the PC control. This loop, adjacent to the previously reported PIP_2_ binding site, showed considerable dynamics, inserting toward the lipid headgroup region upon membrane contact in the Model 1 and PC control simulations (**Figure S2; Movies S2 and S5**) The backbone of the β3-β4 loop in the region of Arg407, Gln408, and Gly409 penetrated at or near the depth of the lipid phosphate plane in each of the simulations (**Table S4 and Figure S6**). The sidechain of Gln408 was at times observed to insert below the phosphate plane in Model 1 and make H-bonds with ester carbonyl oxygens on PC (**Figure S7**).

The only significant penetration observed into the nonpolar portion of the membrane was from the sidechain of Phe452 on the β6-β7 loop, which had an average sidechain insertion depth 2.5 Å below the phosphate plane in Model 1 (**Table S4**). Phe452 inserted near the phosphate plane depth in the other two simulations (**Table S4; Figure S6**).

### Identifying membrane-binding residues via site-directed mutagenesis

In order to test experimentally for the functional importance of the membrane-contacting residues identified in the MD simulations, we selectively mutated residues to alanine, either individually or in combination. Liposome binding activity of each mutant was assessed by measuring the extent of protein-to-membrane FRET for each mutant upon addition of PM, PM(-)PIP_2_, and PM(-)PS/PI liposomes (**Figure 6**). As expected, mutation of residues in the known PIP_2_ binding site (K398A and K410A/K412A) had significant negative effects on binding to PM(-)PS/PI liposomes, in which PIP_2_ is the only target lipid. However, these mutants retained >50% binding toward PM and PM(-)PIP_2_ liposomes, indicating that they are still capable of ample nonspecific electrostatic interactions. The greatest impacts on binding to full PM lipid compositions were found with two triple-mutants: K398A/R451A/R454A and R451A/F452A/R454A; however, even these retained ∼50% of PM liposome binding relative to the wild-type domain (**Figure 6**).

**Figure 6:**
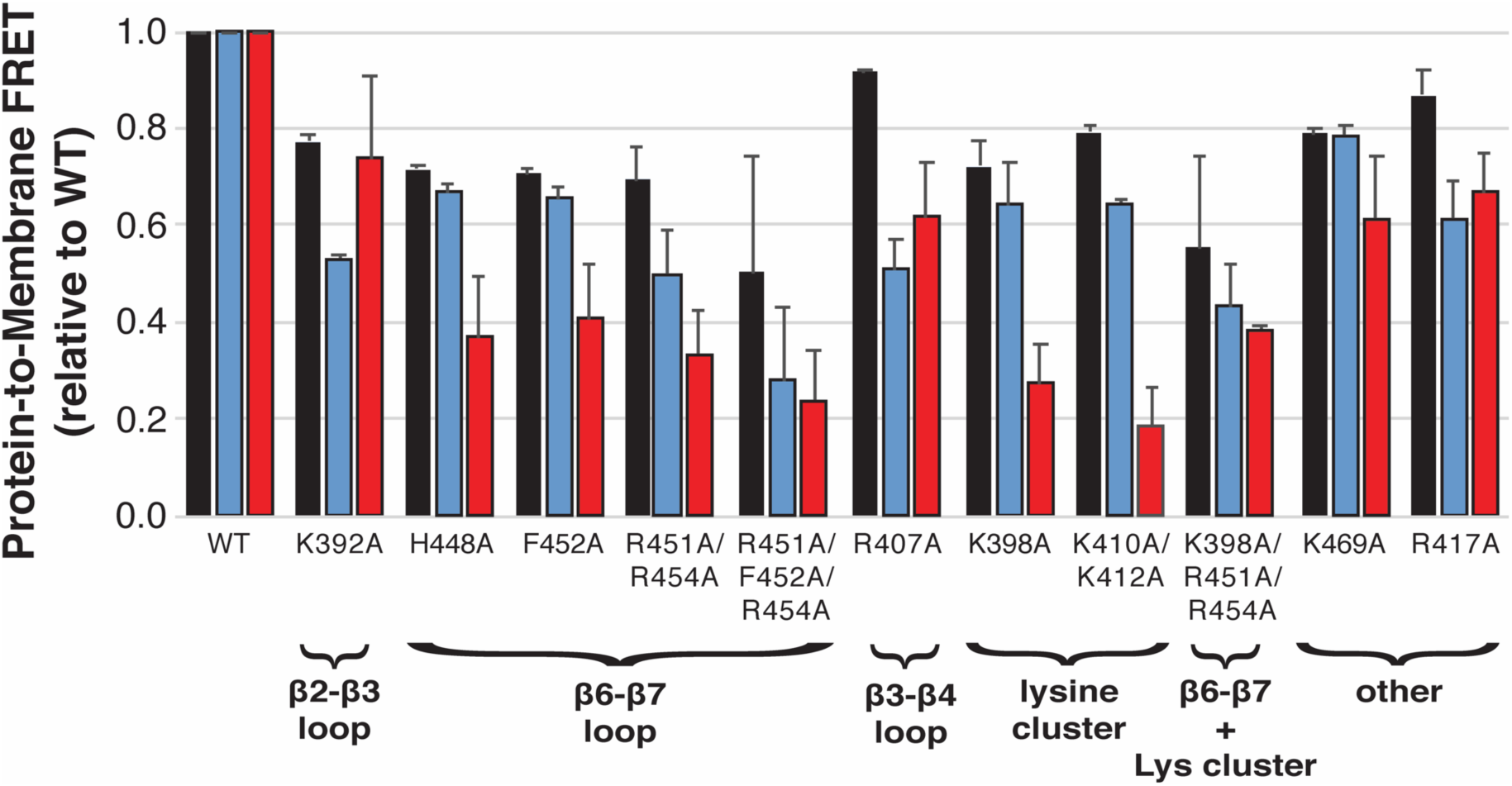
Effects of mutations on Slp-4 C2A liposome interaction. For each indicated mutant C2A domain (1 μM), Trp-to-dansyl FRET was measured upon addition of liposomes (65 μM total accessible lipid) as described in Methods. The extent of FRET was normalized to that of the WT protein domain for each liposome composition. Black, PM; blue, PM(-)PIP_2_: red, PM(-)PS/PI (**Table 1**). Error bars are ± SEM of ≥ 3 replicate samples.

Thermal folding stability of the mutated C2 domains was assessed using differential scanning fluorimetry (**Figure S8**). All of the mutants included in **Figure 6** had clear melting transitions with melting temperatures higher than WT, indicating that neutralization of positive charge and/or mutation of the surface-exposed Phe452 improved the folding stability of the domain. Another mutant, R411A, showed high initial intensity and no melting transition, indicating that the R411A mutant does not fold stably (**Figure S8**). The H448A mutant also had a higher initial intensity, indicating some surface-exposed hydrophobic groups, but retained a clear unfolding transition at a temperature similar to the wild-type domain.

In order to understand how these mutants retain interactions with PIP_2_ and PS, we conducted MD simulations of selected mutant domains binding to PC/PS/PIP_2_ mixed bilayers, using a starting geometry identical to Model 3 of the wild-type protein. The single K398A mutation, in the PIP_2_-binding lysine cluster, resulted in decreased PIP_2_ contact but increased PS contact relative to wild-type, while the single F452A mutation did not change anionic lipid contacts appreciably (**Figure 7**). The triple mutants R451A/F452A/R454A and K398A/R451A/R454A had decreased anionic lipid contact relative to the WT domain (**Figure 7**), particularly in the β6-β7 loop (**Table S5**). Nevertheless, even these triple mutants retained at least 50% of their anionic lipid contacts relative to WT in all simulations, consistent with experimental results (**Figure 6**). For example, loss of Lys398 contact with anionic lipids in some simulations was compensated through increased PIP_2_ contact by Lys410 (**Table S5**). Repeat simulations (varying only by the seed in a random number generator) yielded somewhat different lipid binding contacts for the two triple mutants but did not change the qualitative overall picture, reflecting the availability of a large number of basic residues on the lipid binding surface. Overall, these data from experiments and simulations indicate that multiple mutations decrease membrane affinity somewhat, but the protein retains the ability to bind membranes even when up to three key residues are mutated, consistent with a broad surface of the protein participating in the interaction.

**Figure 7:**
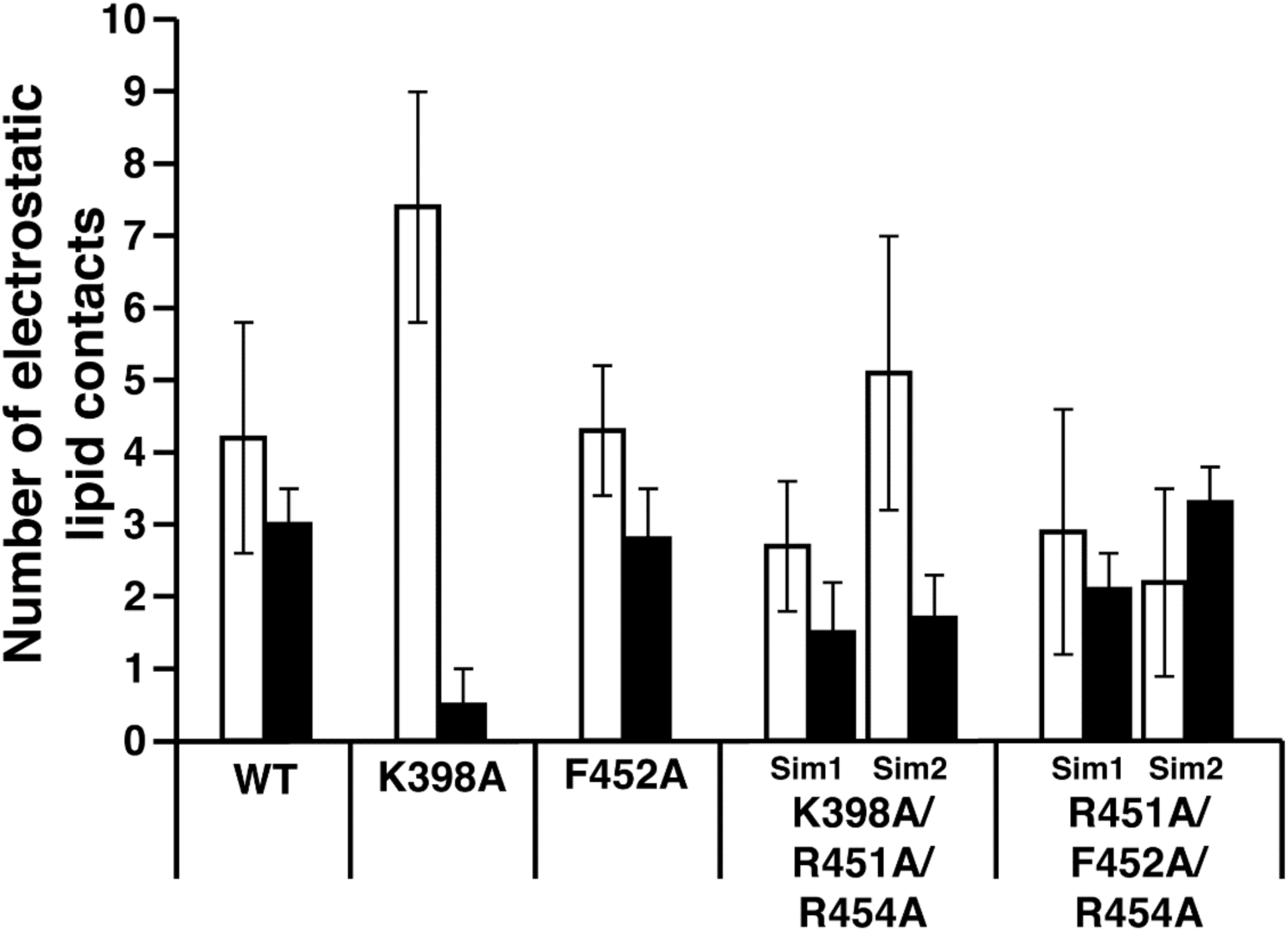
Effects of mutations on Slp-4 C2A lipid binding from MD simulations. The number of basic residues making electrostatic contact with PS (open bars) or PIP_2_ (filled bars) was calculated from the individual residue contacts as described in Methods, averaged (± S.D.) over the final 100 ns of simulation. For the K398A/R451A/R454A and R451A/F452A/R454A mutants, two simulations were conducted which differed only in the random seeds used to calculate initial velocities; data from both simulations for each mutant are shown (Sim1 and Sim2). A breakdown of PS and PIP_2_ contacts by residue for each simulation is given in **Table S5**.

### Identification and characterization of bacterial modifications in the lysine cluster

During the course of protein purification, we noticed that a significant protein peak eluted early during cation exchange chromatography (**Figure 8A**). This is similar to a phenomenon that we and others have reported previously for cationic synaptotagmin C2 domains (22-24). We measured the precise molecular mass of the protein from this peak, and found that the most abundant component had a mass greater than predicted (and compared to the protein in the main peak) by 258 Da (**Figure 8B**). A 258-Da mass increase has been reported previously at or near the N-terminus of bacterially expressed proteins containing an N-terminal His tag, and corresponds to a phosphogluconoyl modification arising from reaction of an amino group on the protein with bacterial phosphogluconolactone (25,26).

**Figure 8:**
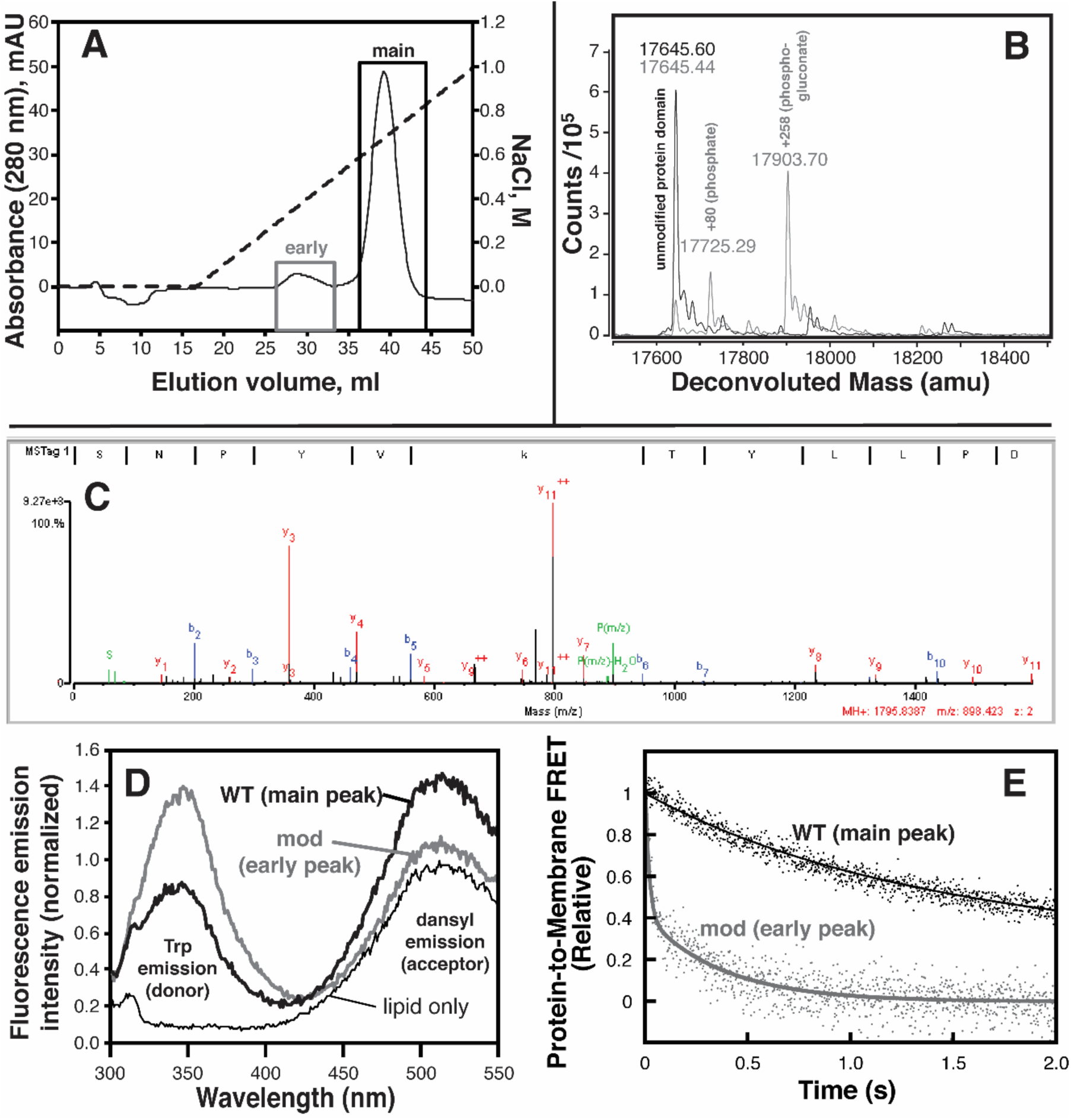
Bacterial post-translational modifications weaken Slp-4 C2A membrane binding. **A**: Chromatogram from cation-exchange protein purification illustrating the two populations of protein collected. Dashed line shows the salt gradient used for elution. **B:** Deconvoluted ESI-QTOF mass spectra of intact protein domains collected from the main (black) and early (gray) peaks in the chromatogram. **C**: Annotated MSMS fragmentation spectrum, including b and y ions, from tandem mass spectrometry of trypsin-digested protein from the early peak. The precursor ion corresponds to the peptide SNPYVkTYLLPD, where k represents phosphogluconoylated lysine at position K398. **D**. Fluorescence emission spectra of PM liposomes before (thin line) and after (thick lines) addition of protein from the early peak (gray) or main peak (black). The emission intensity increase at the dansyl (acceptor) peak is indicative of protein-membrane binding. **E**. Kinetic measurement of dissociation from PM liposomes for the unmodified protein (black, same as **Figure 3**), and modified protein (gray).

As our protein expression system lacks a His tag, we sought to identify the site(s) of modification by performing LC-MSMS experiments on trypsin digests of samples from both peaks. Analysis using proteomics software confirmed the phosphogluconoyl modification on Lys398 in protein from the early peak, for which the fragment ion spectrum contains peaks for the full complement of y-ions as well as most of the b-ions (**Figure 8C)**. In addition, we noted another probable phosphogluconoylation site on Lys412 which we identified via chemical formula matching and manual analysis. Its MSMS fragmentation spectrum includes nearly all predicted y-ions although no b-ions (**Figure S9**). Modification of Lys398 and Lys412 within the lysine cluster must be mutually exclusive, because the whole-protein data only indicate a 258-Da mass increase (**Figure 8B**). Thus, the lysine cluster of the Slp-4 C2A domain is susceptible to modification by reaction with endogenous phosphogluconolactone during bacterial protein expression.

The protein isolated from the early chromatography peak was found to retain a modest binding activity toward PM liposomes, producing a dansyl emission increase that was ∼20% of that of the wild-type protein domain at the same concentration (**Figure 8D**). It is not clear whether this protein-to-membrane FRET arises from weak binding of the phosphogluoconoylated protein, or from another population, e.g. the phosphorylated (+80 Da) population also visible in the whole-domain mass spectra (**Figure 8B**). Due to this relatively low signal, stopped-flow dissociation kinetic data of the modified protein were significantly noisier than wild-type, but could be fit to single- or double-exponential profiles that consistently contained a component with a rate constant of 1.7 ± 0.9 s^-1^ among three independent measurements (two of three measurements also contained a faster component of ∼50 s^-1^) (**Figure 8E**). This is much faster than the major population of the unmodified protein (**Table 3**), consistent with weaker but measurable membrane binding by the modified protein domain. Thus, these data complement our mutational results and show that even a charge-reversal modification does not completely block reversible membrane binding by the Slp-4 C2A domain.

## Discussion

The results presented here show that Ca^2+^-independent binding of the Slp-4 C2A domain to membranes with physiological lipid composition is strong, driven by electrostatics, and robust, its binding activity persisting even when cationic residues are mutated. In particular, we report that (i) physiological levels of PIP_2_ and background anionic lipids (PS and PI) both contribute to a comparable extent toward the thermodynamic stability of the membrane bound state (**Table 2**); (ii) computer modeling and experimental mutational analysis support the lysine cluster as the primary PIP_2_ binding site, whereas basic residues throughout a broad surface contribute to nonspecific anionic lipid binding (**Table 4**); and (iii) the lysine cluster is susceptible to modification by carbonyl compounds such as phosphogluconolactone during bacterial protein expression (**Figure 8**).

### Comparison to other lipid-binding domains

The Ca^2+^-independent nature of Slp-2 C2A contrasts with other well-studied C2 domains, such as those from conventional protein kinase C (PKC) isoforms, synaptotagmin-1, and rabphilins (27-29), and arises due to the absence of key aspartate residues in the β2-β3 and β6-β7 loops (which are called CBL1 and CBL3 in Ca^2+^-sensitive C2 domains). Rather, both C2 domains of Slp-4 lack a full complement of Ca^2+^-coordinating Asp residues but contain a consensus motif associated with PIP_2_ binding, including the lysine cluster in the β3-β4 region (16,30). C2 domains that share these sequence properties include other Slp family C2A domains, Slp-4 C2B, phosphatidylinositol 3-kinase C2A, RIM1 C2B, and synaptotagmin-4 C2A (16,31). This consensus motif is not found in the Ca^2+^-independent C2 domains of PTEN and novel PKC isoforms, which have type II topology (32,33). The mechanism described here is also distinct from the Ca^2+^-independent type I topology C2 domain from kidney and brain-expressed protein (KIBRA), which binds phosphoinositides using the opposite face of the C2 domain (34).

We and others have shown that Slp-4 C2A binds to multiple phosphoinositide species, among which PI(4,5)P_2_ is presumed to be the dominant target due to its ubiquity in the plasma membrane (12,13). The 10-fold enhancement of PIP_2_ affinity in the presence of background anionic lipids is reminiscent of pleckstrin homology (PH) domains that bind phosphoinositides, except that Slp-4 C2A has a measurable affinity for these background anionic lipids even in the absence of PIP_2_ (15,35,36).

### Multiple lipid binding sites

The biexponential dissociation kinetics observed in all of our samples (**Table 3; Table S1**) indicate that the protein exists in more than one lipid binding state. The slowest off rate, ∼0.1 s^-1^, was observed only in lipid compositions that contain both PIP_2_ and background anionic lipids, and therefore most likely represents dissociation from a state with PIP_2_ bound in the lysine cluster and other anionic lipids associated with the broad cationic surface. Faster-dissociating states may then reflect partial occupancy of the nonspecific surface and/or lipids other than PIP_2_ bound in the lysine cluster. The dissociation rate constant of these states tends to decrease (i.e, tighter binding) as more PIP_2_ or PA is present, suggesting that these polyanionic lipids also bind in the nonspecific site(s) (**Table 3; Table S1**).

Binding of PIP_2_ lipids outside of the primary binding site was observed in two of our MD simulations. A PIP_2_ bound to the C-terminal lysine sidechain in Model 1, and two PIP_2_ molecules bound to the β2-β3 and β3-β4 loops, respectively, in Model 2 (**Table 4**). Similarly, binding of phosphoinositide molecules outside of canonical binding sites has been reported computationally and experimentally for the PKCα C2 domain (37), as well as computationally for the GRP-1 PH domain (38).

### Structural insights into membrane binding

Our simulation data indicate that nonspecific anionic lipid interactions include at least 13 basic residues spread over a broad surface of the protein domain (**Table 4, Figure 5B**). These residues include all of the lysine and arginine residues on the lipid-facing surface of the protein except Arg454, which is located on the β6-β7 loop. Its α-carbon was among the closest to the membrane surface in all three PC/PS/PIP_2_ simulations (**Table S4**), and its sidechain projects toward the membrane. Closer inspection revealed that the Arg454 sidechain made extensive contact with the phosphodiester group of PC in all three simulations. Its lack of interaction with PS or PIP_2_ in these simulations could be incidental.

Consistent with a previous report, the K398A mutation effectively blocked binding to PIP_2_ (**Figure 6**, red bars) (17). However, we show that this mutation leaves the protein’s ability to bind background anionic lipids largely intact (**Figure 6**, blue bars). In simulations of WT Slp-4 C2A, Lys398 had little interaction with PS, although it interacted extensively with PIP_2_ in Model 3. Its position in the concave interior of the membrane-binding surface may make it more accessible to large phosphoinositide headgroups than to the smaller headgroups of PS or PA.

Another residue with interesting properties in our study is Arg411. In the crystal structure, this sidechain forms H-bonds with backbone oxygens on the β3-β4 loop and the β5 strand. A role of thse H-bonds in stabilizing secondary structure could explain why the R411A mutant did not fold properly (**Figure S8**). However, in our membrane binding simulations, these H-bonds unraveled as the Arg411 sidechain interacted with lipids. This loss of intra-protein H-bonding likely contributed to the increased conformational flexibility of the β3-β4 loop during membrane binding (**Figure S2)**. It is not yet clear how much this flexibility contributes to the membrane affinity of the domain.

### Significance of phosphogluconoyl modification

We observe that the major protein contaminant in the affinity-purified Slp-4 C2A domain contains a phosphogluconyl modification within the lysine cluster (**Figure 8, Figure S9**). It has been shown previously that certain cationic, PIP_2_-binding C2 domains co-purify with nucleic acids and other contaminants following bacterial expression, and that these contaminants must be removed via ion exchange chromatography in order to properly measure biophysical properties of the C2 domain (22-24). To our knowledge, this report is the first to identify and characterize a particular protein contaminant. We show that the modified protein binds membranes, albeit much more weakly than the unmodified protein (**Figure 8D-E**). This result underscores the importance of the ion exchange step; because the contaminant retains some membrane binding activity, an incompletely purified protein would produce inaccurate results in binding and function assays.

The nature of the modification is surprising: phosphogluconoylation in bacterially purified proteins has been reported previously, but only at the amino terminus of an N-terminal His-tag (25,26,39). In contrast, our expression system used an N-terminal GST tag which was cleaved prior to cation exchange. The location of the modification suggests that the positive electrostatic environment (**Figure 1**) imparts unusually low p*K*_a_ values to the lysine cluster sidechains. This would make the amino groups more efficient nucleophiles for attacking reactive carbonyl compounds, the most abundant of which in *E. coli* BL-21 strains happens to be phosphogluconolactone (25). We speculate that other C2 domains containing lysine clusters likely undergo the same modification.

The physiological significance of the modification is unclear. Although phosphogluconoylation is unlikely to be significant in eukaryotic cells, other carbonyl-containing compounds are known to react nonspecifically with lysines in various diseases including diabetes and alcoholic liver disease (40,41). Indeed, Slp-4 has been identified among the proteins modified by reactive lipid aldehydes in a mouse model of alcoholic fatty liver disease (42). Furthermore, lysines are the target of numerous enzymatic modifications including acetylation and methylation, although it is unknown whether C2 domains are regulated this way. Further work is needed to clarify whether enzymatic or nonenzymatic modification of C2 domain lysine clusters plays a role in their function *in vivo*.

## Experimental and Computational Procedures

### Experimental Materials and Methods

#### Materials

Cholesterol (CH), 1-palmitoyl-2-oleoyl-*sn*-glycero-3-phosphocholine (POPC, PC), 1-palmitoyl-2-oleoyl-*sn*-glycero-3-phosphoethanolamine (POPE, PE), 1-palmitoyl-2-oleoyl-*sn*-glycero-3-phospho-L-serine (POPS, PS), phosphatidylinositol (PI) from liver, phosphatidylinositol-(4,5)-bisphosphate (PIP_2_) from brain, and sphingomyelin (SM) from brain were from Avanti Polar Lipids (Alabaster, AL). N-[5-dimethylamino)-napthalene-1-sulfonyl]-1,2-dipalmitoyl-*sn*-glycero-3-phosphoethanolamine (dansyl-PE) was from NOF America (White Plains, NY). D-*myo*-inositol-(1,2,3,4,5,6)-hexakisphosphate dodecasodium salt (IP_6_) was from Sigma. All reagents were American Chemical Society grade or higher.

#### Protein Cloning, Expression, and Purification

Plasmid DNA encoding human Slp-4 (GenBank ID: BC014913.1) was obtained from American Type Culture Collection (Manassas, VA). The sequence encoding the C2A domain (residues G352-S494) was subcloned into a thrombin-cleavable glutathione-S-transferase fusion vector developed previously for bacterial protein expression and transformed into *E. coli* BL-21 DE3 cells (36,43). Site-directed mutagenesis was performed using the QuikChange II XL kit (Agilent) following the manufacturer’s instructions. All DNA sequences were verified using primer-extension sequencing (Eton Biosciences, San Diego, CA).

Bacterially expressed proteins were purified using glutathione affinity chromatography followed by cation exchange. Cells were lysed in lysis buffer (50 mM Tris, 400 mM NaCl, 1% Triton X-100, 1 mM 2-mercaptoethanol (pH 7.5) with protease inhibitors) by sonication (Sonics VibraCell, Newtown, CT) equipped with a 6-mm probe. Lysates were treated with DNAse I (2 U/mL) from Sigma (St. Louis, MO) for 30 min, centrifuged to remove insoluble matter, and then supernatants were incubated with glutathione sepharose 4B beads (GE Healthcare, Chicago, IL) for 2-3 h at 4 °C. The beads were washed extensively with 50 mM Tris, 400 mM NaCl, 1 mM 2-mercaptoethanol (pH 7.5) and then with 50 mM Tris, 1.1 M NaCl, 5 mM EDTA, 1 mM 2-mercaptoethanol (pH 7.5). Beads were then exchanged into 50 mM Tris, 150 mM NaCl, 0.05 mM EDTA, 1 mM 2-mercaptoethanol (pH 7.7) for cleavage with restriction-grade thrombin (Millipore Sigma, Billerica, MA), and eluted using the thrombin cleavage buffer or buffer A (25 mM HEPES, 15 mM CaCl_2_, 140 mM KCl, 0.5 mM MgCl_2_, pH 7.4) including 1-10 mM 2-mercaptoethanol. Cation exchange chromatography was then performed using an Akta Purifier FPLC system with a HiTrap SP HP column (GE Healthcare), eluting with a gradient of NaCl. Eluted proteins were concentrated using Amicon centrifugal concentrators, purity was assessed using SDS-PAGE, and concentrations were measured based on absorbance at 280 nm using an extinction coefficient of 19060 M^-1^ cm^-1^. Purified proteins were aliquoted, flash-frozen, and stored at -80 °C, and were centrifuged after thawing for 2 min at 17,000 ×g to remove any debris.

#### Liposome Preparation

Phospholipids in chloroform were combined at the desired molar ratio for each experiment (**Table 1**). Lipid films were dried under vacuum for ≥ 2 h and rehydrated in buffer A. Small unilamellar vesicles were prepared by sonication to clarity using a Sonics VibraCell sonicator with a 3-mm tip. Liposomes were stored at 4 °C for at least 8 h after preparation before use, and were used within one week. Lipid concentrations are reported as total accessible lipid, which is approximated as one-half the total lipid.

#### Equilibrium Protein-to-Membrane FRET Measurements

Equilibrium protein-to lipid (Trp-dansyl) FRET titrations were performed as described previously (13). Measurements were made using a Photon Technology International QuantaMaster fluorescence spectrometer at 25 °C, with excitation at 284 nm (1-nm slit width) and emission at 520 nm (8-nm slit width). Protein (1 µM) was pre-mixed with liposomes (125 µM total lipid) in buffer A, and fluorescence was measured upon titration with IP_6_ or NaCl. In a parallel sample, the titrant was added to solutions of lipid alone in order to correct for titrant effects on dansyl fluorescence. For the NaCl titrations, a second correction was made for signal loss upon addition of buffer to protein-lipid mixtures. Samples were equilibrated for 40 s with stirring after each addition. For the IP_6_ titrations, the resulting plot of intensity *F* vs. inhibitor concentration [IP_6_] was fit to a hyperbolic model for single-site competitive inhibition:

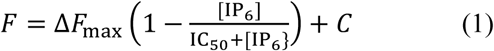

where Δ*F*_max_ is the total FRET signal before ligand titration and IC_50_ is the IP_6_ concentration at which FRET is 50% of the initial value. To simplify graphical representations, data are normalized such that *C* = 0 and Δ*F*_max_ = 1.

In order to compare affinities among membranes of different target lipid compositions, it is both convenient and thermodynamically accurate to model the protein-membrane interaction as partitioning between an aqueous phase and a membrane phase, represented by a mole-fraction partition coefficient which we denote *K*_x_ (44). This thermodynamic constant was calculated from the measured IC_50_ value as follows:

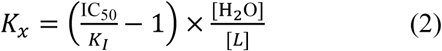

where *K*_I_ is the dissociation constant for protein-IP_6_ binding, reported previously to be 1.8 ± 0.1 µM^-^(13), [H_2_O] is the bulk water concentration, and [*L*] is the total concentration of accessible lipids in the outer leaflet (i.e., half the total bulk lipid concentration). Free energies of binding were then calculated as

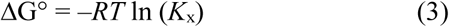

where *R* is the ideal gas constant and *T* is temperature.

For comparison of liposome binding among different mutants, protein-to-membrane FRET measurements were performed using a Cytation 3 fluorescence plate reader (BioTek Instruments, Winooski, VT) with UV-transparent plates (Corning product no. 3679). Tryptophan-to-dansyl FRET was quantified by measuring fluorescence emission of tryptophan in solutions containing the indicated mutant proteins (1 µM, concentrations checked via absorbance immediately prior) in Buffer A with 1 mM TCEP, before and after addition of liposomes (65 µM total accessible lipid). The percentage decrease in tryptophan emission was corrected for intrinsic Trp emission changes upon membrane binding (measured in separate wells using nonfluorescent liposomes) and then normalized to that of the wild-type protein domain.

#### Stopped-Flow Fluorescence Spectroscopy

Stopped-flow fluorescence kinetic measurements were performed using a BioLogic SFM3000 spectrophotometer (Knoxville, TN) using 284 nm excitation and a 455 nm long-pass emission filter. To measure apparent on-rates (*k*_obs_), 1.2 ml of a solution containing protein (0.6 µM) was rapidly mixed with an equal volume of solution containing liposomes (75 µM total accessible lipid) in Buffer A (140 mM KCl, 15 mM NaCl, 0.5 mM MgCl_2_, 25 mM HEPES, 100 µM EDTA, pH 7.4). Protein-to-membrane FRET (dansyl-PE emission) was monitored over time (*t*) for at least 8 replicate shots per sample, which were averaged and fitted to a single-exponenential function (equation 4):

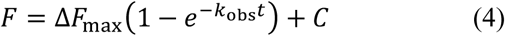

For off rates (*k*_off_), protein-to-membrane FRET (dansyl-PE emission) was monitored following rapid mixing of equal volumes of protein-bound liposomes (75 µM total accessible lipid, 0.6 µM protein) and unlabeled liposomes in Buffer A. Data sets were calculated as the average of 8 or more replicate shots per sample, and were fitted to a single- or double-exponential function (equation 5 or 6, respectively):

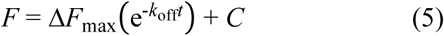

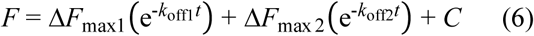

where the *k*_off_ are dissociation rate constants and *C* are offsets.

For simplified presentation, *C* was subtracted and Δ*F*_max_ (or Δ*F*_max1_ + Δ*F*_max2_) normalized to unity in the figures shown. Rate constants listed in **Table 3** are average ± standard deviation of ≥ 3 independent samples. Dead time is estimated to be 1.4 ms.

The reported association rate constants *k*_on,x_ were calculated from the measured *k*_obs_ and *k*_off_ values using the mole-fraction partitioning model (44):

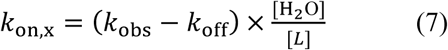

#### Mass spectrometry: Intact Protein Analysis

Purified bacterially expressed and cation exchange separated Slp-4 samples were desalted using C18 ZipTips (EMD Millipore). Desalted samples were diluted with 3% acetonitrile (ACN) in 0.1% formic acid to a final concentation of 100 ng/uL to be analyzed. Samples were chromatographically resolved on-line using a 2.1 x 50 mm, 5.0µ PLRP-S 1000A column (Agilent Technologies) using a 1290 Infinity II LC system (Agilent Technologies). Mobile phases consisted of water + 0.1% formic acid (A) and 90% aq. acetonitrile + 0.1% formic acid (B). Samples were chromatographically separated using a flow rate of 0.3 mL/min using a gradient holding 5% B for 2 minutes and 5-95% B over 3 minutes for a total 5-minute gradient. The gradient method was followed by a column wash at 95% B for 2 minutes before returning to initial condition over 2 minutes. Data was collected on a 6550 Q-TOF equipped with a dual agilent jet stream source (Agilent Technologies) operated in MS-only mode. MS data was collected in positive ion polarity over mass ranges 100– 3200 m/z at a scan rate of 1.5 spectra/sec. Intact protein spectra were deconvoluted using maximum entropy in MassHunter Bioconfirm software (Agilent Technologies) to determine the accurate mass of unmodified and modified protein species if present.

#### Mass spectrometry: Peptide Analysis

Purified Slp-4 samples were tryptically digested using a protein digestion method previously described (45). Samples were reconstituted in 3% acetonitrile + 0.1% formic acid and were chromatographically resolved on-line using a 2.1 x 250 mm, 2.7µ AdvanceBio peptide mapping column (Agilent Technologies) using a 1290 Infinity II LC system (Agilent Technologies). Mobile phases consisted of water + 0.1% formic acid (A) and 90% aq. Acetonitrile + 0.1% formic acid (B). Samples were chromatographically separated using a flow rate of 0.2 mL/min using a gradient holding 5% B over 1 minute, 5-40% B over 9 minutes, and 40-90% B over 2 minutes, for a total 12-minute gradient. The gradient method was followed by a column wash at 90% B for 3 minutes before returning to intial conditions over 2 minutes. Data were collected on a 6550 Q-TOF equipped with a dual agilent jet stream source (Agilent Technologies) operated using intensity-dependent CID MS/MS to generate peptide IDs. MSMS data were collected in positive ion polarity over mass ranges 270–1700 m/z at a scan rate of 10 spectra/s for MS scans and mass ranges 50-1700 m/z at a scan rate of 3 spectra/s for MSMS scans. All charge states were allowed except singly charged species were excluded from being selected during MS/MS acquisition and charge states 2 and 3 were given preference. SpectrumMill software (Agilent Technologies) was used to extract, search, and summarize peptide identity results. Spectra were searched against a custom database containing the Slp-4 protein amino acid sequence allowing up to 2 missed tryptic cleavages with fixed carbamidomethyl (C) and variable deamidated (NQ), oxidation (M) and phosphogluconoyl (K) modifications. The monoisotopic peptide mass tolerance allowed was ± 20.0 ppm and the MS/MS tolerance was ± 50 ppm. A minimum peptide score of 8 and a scored peak intensity of 50% were used as cutoffs for identification of peptides.

### Molecular Dynamics Simulations and Docking Calculations

#### Stand-alone Membrane Model

Two different lipid bilayer membrane models were constructed using the CHARMM-GUI membrane builder (46), each with 256 lipids (128 per leaflet). The first model was made of pure POPC (100 % PC), and the second model had mixed POPC:POPS (190:64, a molar ratio of ∼3:1) with two molecules of 1-stearoyl-2-arachidonoyl-sn-glycero-3-phospho-(D)-myoinositol-(4,5)-bisphosphate (SAPIP_2_, PIP_2_) placed on the protein-proximal leaflet (effective 2% PIP_2_ density). The lipids were described by CHARMM36 force fields with updates for lipids and were hydrated with pre-equilibrated TIP3P (47-49) water models. Randomly selected water molecules were replaced with potassium (K^+^) and chloride (Cl^−^) ions to reach 0.15 M concentration and also for neutralizing the model system. Corrections recommended for the Lennard Jones potential between K^+^ and lipid oxygens were included to avoid unnaturally strong binding of K^+^ to anionic lipids (50).

Each solvated model was minimized for 21,000 steps followed by equilibration for 2.8 ns at 1 bar and 298 K using NAMD version 2.10 (51). The production run of equilibration was then extended for 200 ns under the same conditions. Temperature was controlled with Langevin dynamics where the temperature dampening coefficient was set to 1.0 and pressure controlled using the Langevin piston method (52,53) with an oscillation period of 75 fs and a damping time scale of 25 fs. Long-range electrostatic interactions were computed using the particle mesh Ewald method (54,55), and the short-range non-bonded interactions cut-off was set to 12 Å with a switching function set to 11 Å. The SHAKE algorithm was used to make waters rigid as well as to constrain all bonds between hydrogens and heavy atoms (56). A time step size of 2 fs was adopted. Unless otherwise stated, the same parameters were applied in the dynamics simulations throughout this paper. The constructed membrane models were validated by examining the area per lipid (APL) and order parameters (*S*_CH_) (57), which were found to be close to available literature values (50,58-60) (**see Additional Details of Methods in the Supporting Information**).

#### Stand-alone Protein Model

The crystal structure of the Slp-4 C2A domain (PDB: 3FDW) with all hydrogens added was solvated in the TIP3P water box at a physiological salt concentration (0.15 M KCl) and pH 7. The protein was described by the CHARMM36 force fields. The model was minimized for 10,000 steps with the protein backbone atoms (N, Cα, C, and O) frozen. Then, the system was slowly heated from 0 to 298 K for 40 ns with the backbone frozen. The model was further equilibrated for 2 ns at 298 K without constraints. Finally, an extended production run was performed for 200 ns. The trajectory was saved every 100 ps. Snapshots extracted from the saved trajectory were used in subsequent docking calculations and in the building of protein-membrane complex models. The electrostatic potential maps (+1.0 kT/e in blue and −1.0 kT/e in red, electrostatic equipotential contours) were calculated using APBS-PDB2PQR (61,62) at pH 7.0 with 0.15 M KCl.

The protein exhibited certain conformational changes involving mostly the following regions: (i) A386 to K390 of the β2-β3 loop, (ii) H449 to N455 of the β6-β7 loop, (iii) P403 to G409 of the β3-β4 loop, and (iv) K398/K410/K412 which comprise the lysine cluster. To describe the conformation of the protein, we defined two angles using the tips of all three defined loops (β2-β3, β6-β7, and β3-β4) and the center-of-mass (COM) of the protein (**Figure S2A)**. The tip of each loop was defined as the center of mass of the C_α_ of three consecutive residues. More specifically, the tip of β2-β3 was represented by E388, A389, and K390; the tip of β6-β7 by G450, R451, and F452; and the tip of β3-β4 by S406, R407, and Q408. For example, in the crystal structure, the angle between β2-β3 and β6-β7 (which we term angle *α*) is 35°, and the adjacent angle between β6-β7 and β3-β4 (which we term angle *β*) is 76°.

#### Docking Calculations

To determine the location of PIP_2_ binding to the surface of the Slp-4 C2A domain, docking calculations were performed with inositol-(3,4,5)-trisphosphate (IP_3_, the soluble analogue of the PIP_2_ head group) as the ligand and the entire C2A domain (either wild-type or mutant) as the receptor using the Flexidock module in Sybyl 8.0. For the WT protein, we employed the experimental protein structure (PDB: 3FDW) as well as three equilibrated protein structures extracted from the simulated trajectory of the stand-alone protein at *t* = 9.2, 120, and 200 ns showing different protein conformations. *In silico* mutations were performed on the two equilibrated WT protein structures (*t* = 9.2 and 200 ns) using the Sybyl 8.0 mutation tool, followed by geometry minimizations. The ligand and proteins were described by the Tripos force fields (63). The Kollman (64) all-atom charges were assigned to the protein and Gasteiger-Huckel (65,66) charges to the ligands. Rotatable bonds were allowed in the ligand but not in the protein. The docked poses were clustered (**see Additional Details of Methods in the Supporting Data**) and the largest cluster can be interpreted as the most probable location of IP_3_ binding.

#### Protein-Membrane Complexes

We first built the complex model for the WT protein and membranes. Briefly, the last snapshot of the trajectory (i.e. *t* = 200 ns) of the equilibrated stand-alone WT protein and membrane structures were merged using CHARMM (c40b1) and VMD (67,68). We constructed four protein-membrane complex models: a PC Control featured the POPC membrane model and Models 1–3 the mixed PC/PS/PIP_2_ membrane model (**Figure S3**). In all models, the protein was placed above the membrane with the protein COM ∼21–25 Å above the average phosphate (PO_4_) plane of the upper leaflet of the lipid bilayer. In Models 1–3, the proteins faced the membrane surface differently: (i) Model 1 had the β2-β3 and β6-β7 loops positioned between the two PIP_2_ lipids such that each PIP_2_ lipid was of approximately equal distance from the both loops, distances of 25-30 Å. (ii) Model 2 was made by rotating the protein in Model 1 approximately 90° counterclockwise about the *z-*axis. The distance between one PIP_2_ and the loops region was about the same as the distance between the other PIP_2_ and the β4 binding site (∼15–20 Å). (iii) In Model 3, β4 in the protein was positioned directly above one of the PIP_2_ lipids. Each model was minimized for 21,000 steps and equilibrated for 2.8 ns, followed by a production run for 200 ns using NAMD.

Next, we constructed the complex models for two single mutants K398A and F452A and two triple mutants R451A/F452A/R454A and K398A/R451A/R454A using the same protocol as for WT except for some minor differences. A mutant complex model employed the mixed PC/PS/PIP_2_ membrane model and a starting protein geometry identical to Model 3 of the WT protein. The selected residue(s) was(were) first mutated *in silico* and minimized while keeping the rest of the protein frozen. The numbers of K^+^ and Cl^−^ ions were adjusted to keep the system charge neutral and at 0.15 M salt concentration. The mutant complex model was then minimimized for 10,000 steps and equilibrated for 5 ns, followed by production runs in the same way as for the WT complex models.

#### Depth Penetration Calculations

Depth penetration calculations were carried out to highlight penetration that occurred in the targeted regions. The position of the membrane was defined as the average *z*-coordinate of all the phosphorus atoms in the protein-facing leaflet of the bilayer. The selected residues were represented as the depth of their C_α_ atoms, except where noted. The membrane plane was set as the reference for the depth penetration calculated as the difference between the *z*-coordinates of the C_α_ of the given residue and of the membrane plane, with more negative values corresponding to deeper insertion.

#### Electrostatic Contacts

Electrostatic contacts were defined as the selected basic residue sidechains being within a cutoff distance from the heavy atoms of anionic lipid headgroups (from the phosphodiester linkage outward) in the model systems containing PS and PIP_2_. Calculations were performed for arginine and lysine as well as for histidine (H376, H476, and H481 as the HSE tautomer; H381, H448, and H449 as HSD), which was uncharged in all simulations. Cutoff distances were as follows: 5 Å for the amine nitrogen (N_𝓏_) of lysine, 6.3 Å for the guadinino carbon (C_𝓏_) of arginine, and 6.1 Å for the imizadole ring COM of histidine. For the PC control, similar calculations were performed for contacts with PC headgroups, considering heavy atoms from the glycerol backbone outward. Electrostatic lipid contacts were calculated independently for each residue; if a given residue was within the cutoff distance of one or more atoms of a particular lipid molecule, this was counted as a single contact.

## Author Contributions

A.A., A.W-S., C.T.S., H.L., and J.D.K. designed and conceived experiments; N.L.C., H.L., and J.D.K. designed and conceived simulations; A.A., A.W.S, M.N., J.R.O., J.O., T.L., C.M., and J.D.K. performed experiments; S.T., M.N., J.A.H., and N.L.C. set up and conducted simulations; A.A., A.W-S., S.T., M.N., J.A.H., J.R.O., N.L.C., J.O., T.L., C.M., R.R., H.L., and J.D.K. analyzed data; A.A., S.T., M.N., J.A.H., N.L.C., C.M., N.R., R.R., C.T.S., H.L, and J.D.K. wrote the paper.

