## Supporting Information for "Multivalent lipid targeting by the calcium-independent C2A domain of Slp-4/granuphilin"

#### Additional Details of Methods

##### Assessment of Membrane Models

To assess the constructed membrane models, area per lipid (APL) and order parameters ($S_{CH}$) were calculated. The APL was calculated by dividing the total area of lipid by the total number of lipids of each leaflet (128 lipids); the total area was averaged over the 200 ns of the productive run. The APL for pure POPC, 0.64 $nm^2$, matches closely to the reference APL 0.639 $nm^2$. (1,2) The $S_{CH}$ describes the orientation of the lipid alkyl chains. The value of $S_{CH}$ is determined by measuring the angle $\theta$ between the vector normal to the membrane surface and the vector of C-H bond of the lipid (Eq. S1).[14]

$$S_{\mathrm{CH}} = \frac{1}{2}(3\langle\cos^2\theta\rangle - 1) \qquad \text{(S1)}$$

The brackets indicate the average over the two C-H bonds in a $CH_2$ group at a given position along the alkyl chain over all lipids of the same type and over the 200 ns of simulation time. The $S_{CH}$ has yet to be determined for the arachidonic and stearic acyl chains of $PIP_2$. However, reference values for the oleoyl and palmitoyl acyl chain have been determined, and are in close agreement with $S_{CH}$ values found in this work. (3,4) We therefore conclude that the membrane models are valid for use in the subsequent simulations.

##### Docking Calculations and Clustering

For each protein conformation tested, 510 different initial positions of the ligand were distributed uniformly around the protein. The 510 docked poses were clustered based on the distances between the centers of mass of the ligands, using a cutoff distance of 10 Å. More specifically, we began with the pose of the highest fitness, which was the delegate of the first cluster. We then assigned to the first cluster the docked ligands whose centers of mass were within 10 Å of the center of mass of the first-cluster delegate. All poses in the first cluster were deleted from the list of 510 poses. Next, we identified the pose of the highest fitness among the remaining poses, made it the delegate of the second cluster, and included poses within 10 Å of the center of mass of this delegate into the second cluster. This process was repeated until every pose was assigned to a cluster. In this way, the clusters are mutually exclusive, because every pose can only be assigned to one cluster.

**Figure S1**

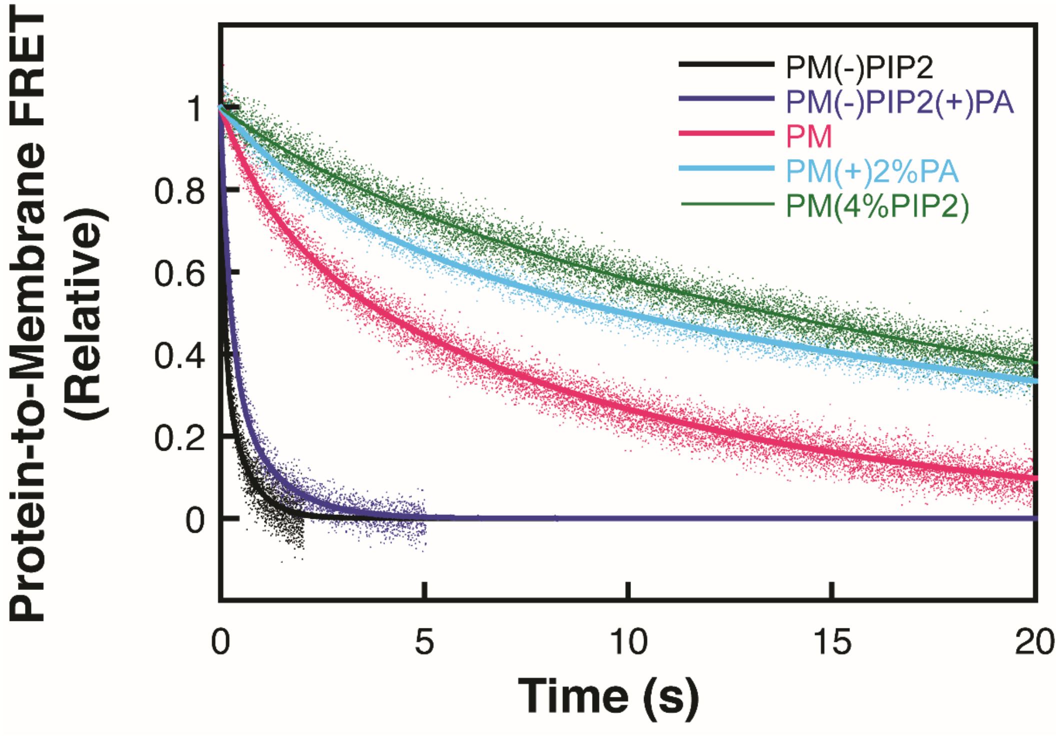

**Figure S1: Effect of phosphatidic acid (PA) on wild-type Slp-4 C2A membrane dissociation kinetics.** Kinetics were measured using stopped-flow fluorescence spectroscopy and fit as described in the Methods of the main text. Fit parameters are given in **Table S1**. PA has a small headgroup with a net charge of approximately -1.5 at pH 7.4. (5) Inclusion of 2% POPA in the liposome lipid composition decreased the off-rate by approximately a factor of two, both in the absence (compare dark blue to black curves) and in the presence (compare cyan to magenta curves) of 2% $PIP_2$. For comparison, inclusion of 2% $PIP_2$ decreased the off-rate by a factor of approximately 40 (compare magenta to black curves), but a further increase to 4% $PIP_2$ only decreased the off-rate further by a factor of approximately two (compare green to magenta curves). These results suggest that despite its polyanionic character, PA cannot substitute for $PIP_2$ in the PIP-selective binding site of the C2 domain, but rather slows protein dissociation through nonspecific secondary electrostatic interactions. The results are consistent with $PIP_2$ binding the protein both selectively (dominant at lower $PIP_2$ percentages) and and through nonspecific electrostatic interactions (apparent at higher $PIP_2$ percentages).

Lipid compositions are based on those shown in **Table 1** of the main text; PA was substituted for PC, whereas $PIP_2$ was substituted for PI.

**Table S1: Comparison of rate constants ($k_1$, $k_2$), amplitudes ($Amp_1$, $Amp_2$), and time to 50% dissociation ($t_{1/2}$) from the data shown in Figure S1.** Two independent sets of measurements were performed and both values are shown here.

| Lipid composition | Lipid comp alt | $k_1$ ($s^{-1}$) | $Amp_1$ (%) | $k_2$ ($s^{-1}$) | $Amp_2$ (%) | $t_{1/2}$ (s) |
|---|---|---|---|---|---|---|
| PM(-)PIP2 | 0% PIP2 0% PA | 12, 10 | 60, 100 | 1.8, ND | 40, 0 | 0.11, 0.07 |
| PM(-)PIP2(+)PA | 0% PIP2 2% PA | 4.3, 8.0 | 62, 64 | 0.94, 1.0 | 38, 36 | 0.26, 0.17 |
| PM | 2% PIP2 0% PA | 0.72, 1.35 | 28, 22 | 0.10, 0.17 | 72, 72 | 4.0, 2.7 |
| PM(+)2%PA | 2% PIP2 2% PA | 0.31, 0.70 | 30, 35 | 0.037, 0.079 | 70, 65 | 9.9, 3.9 |
| PM(4%PIP2) | 4% PIP2 0% PA | 0.30, 0.42 | 12, 20 | 0.042, 0.053 | 88, 80 | 13.6, 9.0 |

**Figure S2**

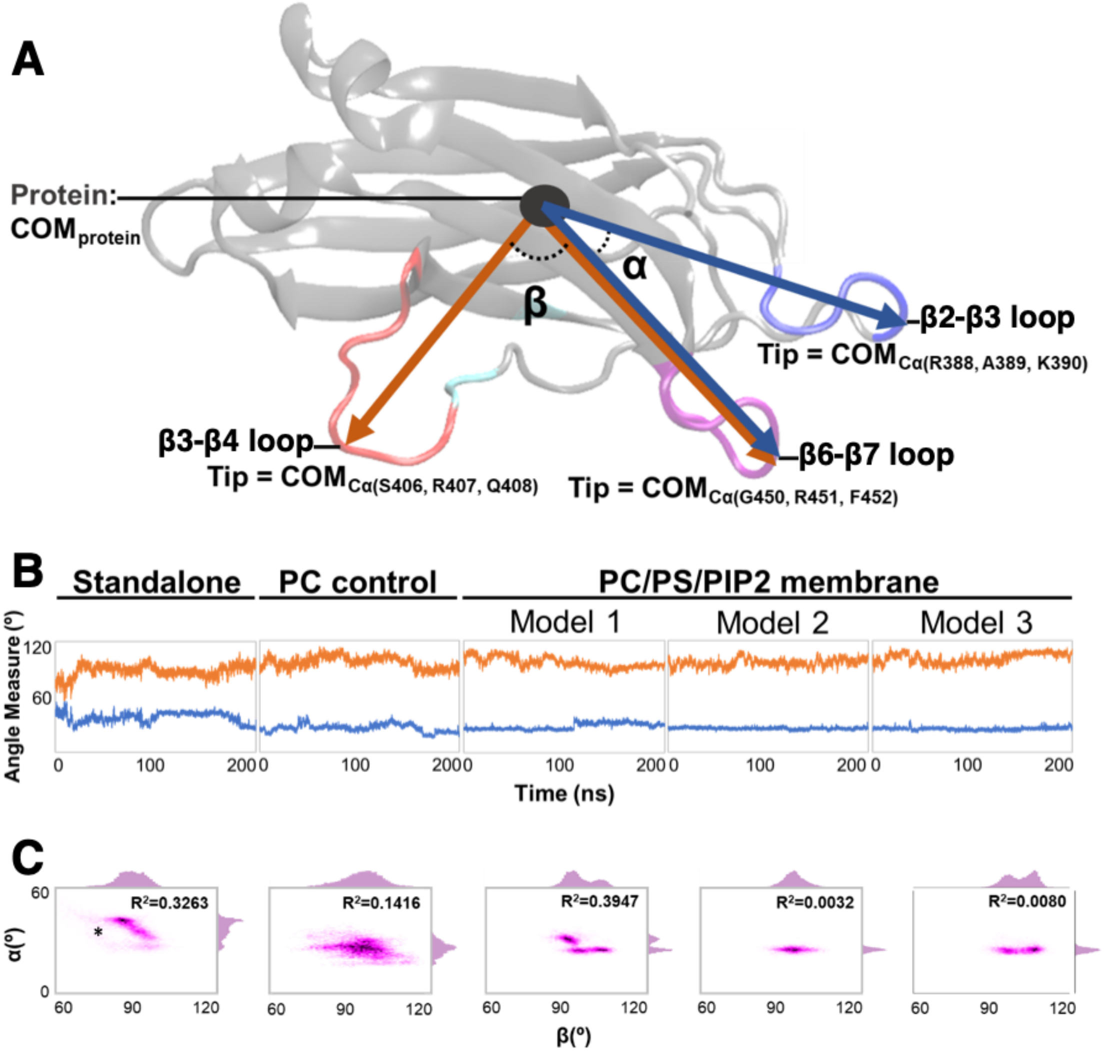

**Figure S2: Protein conformational changes during MD simulations.** **A:** Structure of the Slp-4 C2A domain illustrating the definitions of angles α and β which we use to report protein conformational changes. **B:** Angle values of α (blue) and β (orange) during MD simulations of the protein alone (left), or initially positioned above a lipid bilayer. **C:** Correlation plots of α and β during each simulation. Asterisk indicates α (35°) and β (76°) values in the crystal structure.

During the course of the standalone protein simulation, α decreased while β increased in a correlated manner, indicating movement of the β6-β7 loop toward the β2-β3 loop. The final conformation from the standalone simulation was used as the starting protein conformation in each of the simulations with lipids. Although α and β exhibited broad, uncorrelated dynamics in the PC control simulation, the simulations in the presence of anionic membranes confined α to a narrow range of values with the β6-β7 and β2-β3 loops in close proximity. Correlation analysis suggests the presence of up to three distinct conformational substates in these simulations. The greater variability of β reflects dynamics of the β3-β4 loop.

**Table S2: Results of $IP_3$ docking calculations toward mutant forms of Slp-4 C2A.** Docking calculations were performed using FlexiDock as described in the main text. The lysine cluster and loops cluster are defined as in **Figure 4A** of the main text. Mutations were generated computationally using the indicated starting structure from the standalone simulation of the wild-type protein domain.

| Protein Structure | Snapshot 1 (9.2 ns) (α = 55°) | | | Snapshot 3 (200 ns) (α = 25°) | | |
|---|---|---|---|---|---|---|
| | Lys cluster (%) | Loops (%) | Other (%) | Lys cluster (%) | Loops (%) | Other (%) |
| **Wild Type** | 81 | 5 | 14 | 33 | 32 | 35 |
| **K398A** | 65 | 10 | 25 | 12 | 42 | 45 |
| **K410A** | 68 | 7 | 25 | 12 | 39 | 48 |
| **K412A** | 69 | 12 | 18 | 8 | 48 | 45 |
| **K398A/K410A** | 53 | 13 | 29 | 4 | 42 | 54 |
| **K398A/K412A** | 46 | 21 | 33 | 0 | 44 | 57 |
| **K410A/K412A** | 33 | 27 | 40 | 0 | 47 | 54 |
| **K398A/K410A/K412A** | 9 | 45 | 47 | 0 | 45 | 55 |

#### Figure S3

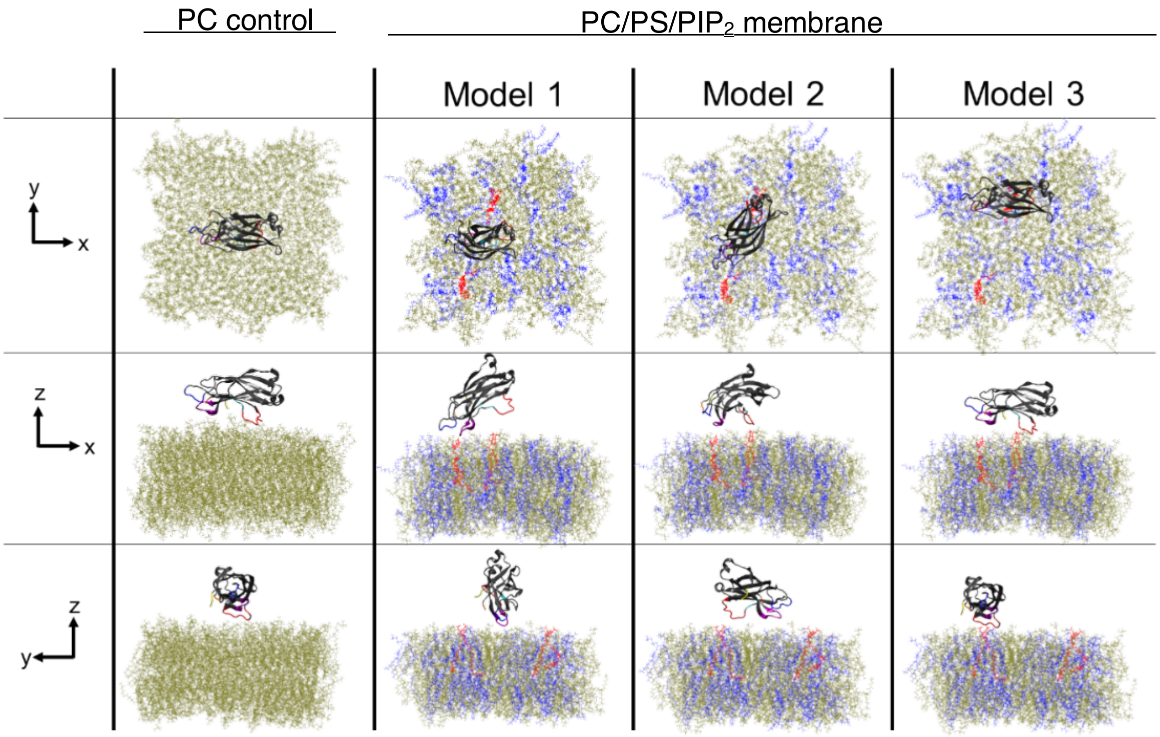

**Figure S3: Initial geometries used for MD simulations with membranes.** Each model comprises a protein membrane system (solvated in water) that was run for 200 ns. All models were solvated in 0.15 M KCl to maintain a physiological salt concentration. The initial protein conformation used for all simulations is the endpoint (200 ns) of the protein standalone simulation. In Models 1, 2, and 3 the protein domain (gray) sits on top of a membrane bilayer that is made up of 190 PC molecules (tan), 64 PS molecules (blue), and 2 $PIP_2$ molecules (red). These models differ in the initial placement of the protein domain and lipids. In Model 1, the protein domain was placed between the two $PIP_2$ molecules with its lysine cluster toward the membrane and its long axis approximately perpendicular to a line joining the $PIP_2$ molecules. In Model 2, the long axis was approximately parallel to a line joining the $PIP_2$ molecules; i.e., the two $PIP_2$ molecules were underneath the β2-β3 and β3-β4 loops, respectively. In Model 3, the protein was placed with its lysine cluster directly above one of the $PIP_2$ molecules. In the PC control, the protein was placed centrally with its initial orientation the same as Model 3.

**Table S3:** Full list of electrostatic contacts between basic residues and anionic lipids in the PC/PS/$PIP_2$ simulations. All basic residues are included. The number of lipid molecules contacted by each residue was averaged over the last 100 ns of simulations (± S.D.). Contacts were calculated as described in Methods. Secondary structural elements are listed on the left, and the conserved lysine cluster is highlighted blue.

| | | Model 1 | | Model 2 | | Model 3 | |
|---|---|---|---|---|---|---|---|
| | | Avg PS Contact | Avg $PIP_2$ Contact | Avg PS Contact | Avg $PIP_2$ Contact | Avg PS Contact | Avg $PIP_2$ Contact |
| β1 | **R359** | 0.00 ± 0.00 | 0.00 ± 0.00 | 0.00 ± 0.00 | 0.00 ± 0.00 | 0.00 ± 0.00 | 0.00 ± 0.00 |
| | **K365** | 0.00 ± 0.00 | 0.00 ± 0.00 | 0.01 ± 0.09 | 0.00 ± 0.00 | 0.00 ± 0.00 | 0.00 ± 0.00 |
| β2 | **H376** | 0.00 ± 0.00 | 0.00 ± 0.00 | 0.00 ± 0.00 | 0.00 ± 0.00 | 0.00 ± 0.00 | 0.00 ± 0.00 |
| | **K378** | 0.00 ± 0.00 | 0.00 ± 0.00 | 0.00 ± 0.00 | 0.00 ± 0.00 | 0.00 ± 0.00 | 0.00 ± 0.00 |
| | **H381** | 0.00 ± 0.00 | 0.00 ± 0.00 | 0.00 ± 0.00 | 0.00 ± 0.00 | 0.00 ± 0.00 | 0.00 ± 0.00 |
| β2- β3 | **K390** | 1.30 ± 0.79 | 0.00 ± 0.00 | 0.30 ± 0.45 | 0.10 ± 0.04 | 0.01 ± 0.11 | 0.00 ± 0.00 |
| | **K391** | 0.00 ± 0.01 | 0.00 ± 0.00 | 0.00 ± 0.00 | 0.00 ± 0.00 | 0.00 ± 0.00 | 0.00 ± 0.00 |
| | **R392** | 1.12 ± 0.01 | 0.00 ± 0.00 | 0.13 ± 0.34 | 1.00 ± 0.00 | 0.45 ± 0.69 | 0.00 ± 0.00 |
| β3 | **K398** | 0.00 ± 0.04 | 0.00 ± 0.00 | 0.00 ± 0.00 | 0.00 ± 0.00 | 0.59 ± 0.49 | 1.00 ± 0.06 |
| β3-β4 | **K405** | 0.00 ± 0.05 | 0.00 ± 0.00 | 0.01 ± 0.11 | 0.99 ± 0.10 | 0.33 ± 0.48 | 0.00 ± 0.00 |
| | **R407** | 0.08 ± 0.27 | 0.00 ± 0.00 | 0.19 ± 0.39 | 1.00 ± 0.00 | 0.90 ± 0.45 | 0.00 ± 0.00 |
| | **K410** | 0.00 ± 0.00 | 0.00 ± 0.00 | 0.00 ± 0.00 | 0.00 ± 0.00 | 0.37 ± 0.50 | 0.00 ± 0.00 |
| β4 | **R411** | 1.59 ± 0.60 | 0.00 ± 0.00 | 0.19 ± 0.39 | 0.00 ± 0.00 | 0.65 ± 0.48 | 0.00 ± 0.00 |
| | **K412** | 0.78 ± 0.54 | 0.00 ± 0.00 | 0.12 ± 0.33 | 0.00 ± 0.00 | 0.02 ± 0.13 | 1.00 ± 0.00 |
| β4-β5 | **K416** | 0.29 ± 0.51 | 0.00 ± 0.00 | 0.52 ± 0.52 | 0.00 ± 0.00 | 0.00 ± 0.04 | 0.00 ± 0.00 |
| | **R417** | 0.24 ± 0.45 | 0.02 ± 0.13 | 0.97 ± 0.55 | 0.10 ± 0.31 | 0.26 ± 0.49 | 0.01 ± 0.08 |
| β5 | **R429** | 0.09 ± 0.29 | 0.00 ± 0.00 | 0.26 ± 0.49 | 0.00 ± 0.00 | 0.01 ± 0.10 | 0.00 ± 0.00 |
| β6 | **R440** | 0.00 ± 0.00 | 0.00 ± 0.00 | 0.00 ± 0.00 | 0.00 ± 0.00 | 0.00 ± 0.04 | 0.00 ± 0.00 |
| | **H448** | 0.00 ± 0.00 | 0.00 ± 0.00 | 0.00 ± 0.00 | 0.00 ± 0.00 | 0.00 ± 0.00 | 0.00 ± 0.00 |
| β6-β7 | **H449** | 0.02 ± 0.13 | 0.00 ± 0.00 | 0.17 ± 0.39 | 0.00 ± 0.00 | 0.01 ± 0.07 | 0.91 ± 0.29 |
| | **R451** | 0.00 ± 0.00 | 0.00 ± 0.00 | 0.52 ± 0.50 | 0.00 ± 0.06 | 0.51 ± 0.80 | 0.00 ± 0.00 |
| | **R454** | 0.00 ± 0.00 | 0.00 ± 0.00 | 0.00 ± 0.00 | 0.00 ± 0.00 | 0.12 ± 0.32 | 0.12 ± 0.33 |
| β7-β8 | **K469** | 0.00 ± 0.00 | 0.00 ± 0.00 | 0.00 ± 0.00 | 0.00 ± 0.00 | 0.00 ± 0.00 | 0.00 ± 0.00 |
| | **K472** | 0.00 ± 0.00 | 0.00 ± 0.00 | 0.00 ± 0.00 | 0.00 ± 0.00 | 0.00 ± 0.00 | 0.00 ± 0.00 |
| | **K473** | 0.00 ± 0.00 | 0.00 ± 0.00 | 0.00 ± 0.00 | 0.00 ± 0.00 | 0.00 ± 0.00 | 0.00 ± 0.00 |
| β8 | **H476** | 0.00 ± 0.00 | 0.00 ± 0.00 | 0.00 ± 0.00 | 0.00 ± 0.00 | 0.00 ± 0.00 | 0.00 ± 0.00 |
| C-ter | **H481** | 0.00 ± 0.00 | 0.00 ± 0.00 | 0.02 ± 0.15 | 0.00 ± 0.00 | 0.00 ± 0.00 | 0.00 ± 0.00 |
| | **K483** | 0.00 ± 0.00 | 0.95 ± 0.21 | 0.00 ± 0.00 | 0.00 ± 0.00 | 0.00 ± 0.04 | 0.00 ± 0.00 |

**Figure S4**

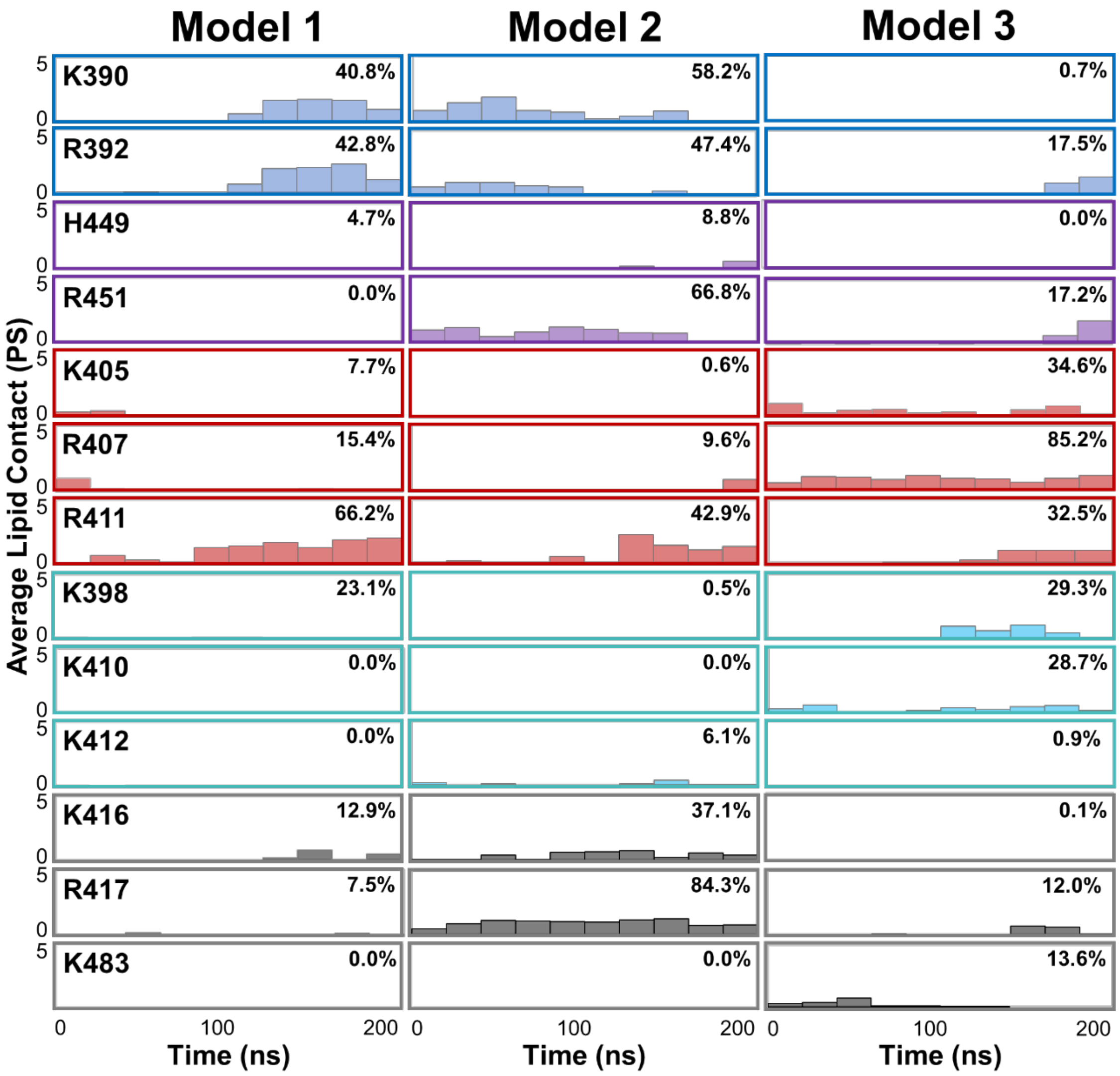

**Figure S4. PS electrostatic contacts.** Bars represent the POPS contact number (averaged over 20-ns intervals) for the thirteen residues with the most total electrostatic contact throughout the 200-ns simulations. The percentage in the top right of each graph represents the proportion of the simulation during which at least one contact was observed. Data sets are colored by structural element: β2-β3 loop (blue), β6-β7 loop (purple), β3-β4 loop (red), the conserved lysine cluster (cyan), and other residues in other regions of the protein (gray). Interactions with the β2-β3 loop were lowest in Model 3, while interactions with the β6-β7 and β3-β4 loops varied throughout the models. PS binding in addition to $PIP_2$ was detected in the lysine cluster site in Model 3. Arg411 interaction with PS increased over time in all of the simulations.

#### Figure S5

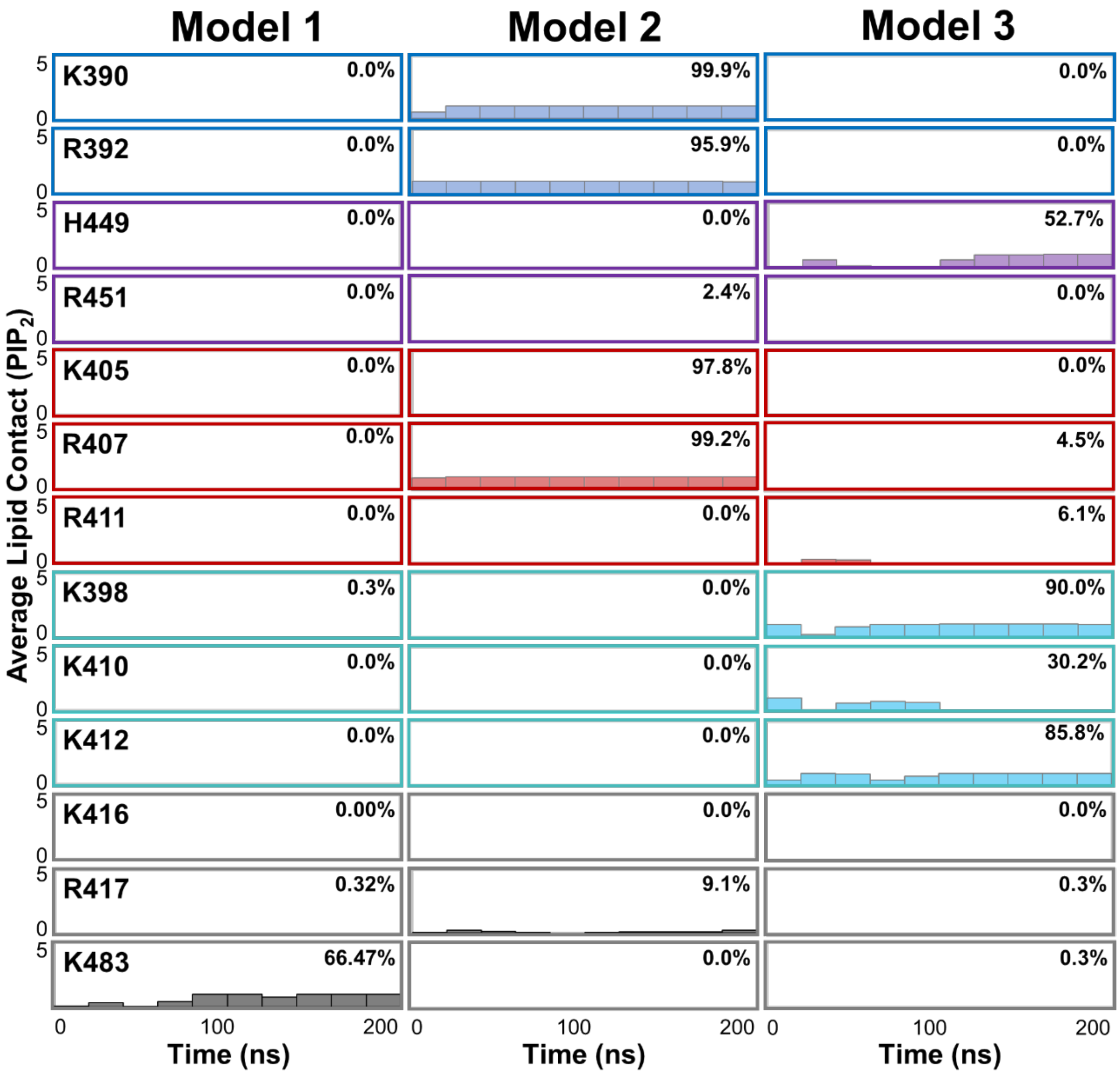

**Figure S5. $PIP_2$ electrostatic hydrophilic contacts.** Bars represent the $PIP_2$ contact number (averaged over 20-ns intervals) for the thirteen residues with the most total electrostatic contact throughout the 200-ns simulations. The percentage in the top right of each graph represents the proportion of the simulation during which at least one contact with $PIP_2$ was observed. Data sets are colored by structural element: β2-β3 loop (blue), β6-β7 loop (purple), β3-β4 loop (red), the conserved lysine cluster (cyan), and residues in other regions of the protein (gray). In all three models, $PIP_2$ interacted with the residues directly above it in the initial placement: K483 (C-terminus) in Model 1, the β2-β3 and β3-β4 loops in Model 2, and the lysine cluster plus H449 in Model 3. The contacts in Model 3 most closely resemble the previously reported binding surface for this domain (6) as well as other $PIP_2$-binding C2 domains. (7-9)

**Table S4: Penetration depth (Å) of selected residues during MD simulations.** Distances relative to the average phosphate plane of the membrane are presented for the given residues in the PC control and Models 1, 2, and 3. Data are presented as mean ± standard deviation (minimum/maximum) over the 200-ns simulation. All values shown are from backbone α-carbons, unless otherwise noted. Overall, most of the protein-lipid interactions are with the polar headgroup region (positive values) rather than below the phosphate plane (negative values).

| | | PC control | PC/PS/PIP2 membrane | | |
|---|---|---|---|---|---|
| | | | Model 1 | Model 2 | Model 3 |
| Protein | | 26.0 ± 2.4 (21.1 / 34.7) | 14.7 ± 2.4 (10.3 / 20.9) | 15.9 ±1.2 (12.8 / 20.5) | 18.1 ± 2.5 (12.7 / 24.5) |
| β2-β3 loop | A386 | 28.2 ± 4.6 (15.5 / 39.0) | 8.7 ± 1.5 (3.2 / 13.3) | 12.6 ± 2.3 (6.7 / 19.3) | 12.4 ± 3.2 (4.7 / 23.2) |
| | D387 | 28.0 ± 4.2 (17.7 / 39.6) | 7.6 ± 1.7 (1.3 / 12.4) | 10.9 ± 2.2 (5.2 / 17.9) | 11.9 ± 3.7 (4.1 / 24.5) |
| | E388 | 30.0 ± 4.8 (19.4 / 42.9) | 10.4 ± 2.0 (2.1 / 15.7) | 11.3 ± 2.4 (7.5 / 20.2) | 14.1 ± 3.8 (5.8 / 27.5) |
| | A389 | 32.7 ± 5.1 (21.2 / 46.0) | 8.8 ± 2.1 (0.3 / 14.8) | 11.3 ± 2.4 (5.3 / 19.2) | 13.6 ± 4.2 (4.8 / 28.6) |
| | K390 | 32.8 ± 5.7 (18.7 / 45.9) | 7.7 ± 1.7 (2.3 / 13.4) | 11.8 ± 2.0 (5.6 / 19.0) | 16.2 ± 4.9 (7.0 / 31.6) |
| β3-β4 loop | Y400 | 19.8 ± 2.4 (15.1 27.8) | 14.7 ± 2.7 (6.4 / 18.2) | 13.9 ± 1.2 (10.4 / 18.3) | 13.7 ± 2.3 (8.4 / 21.4) |
| | L401 | 19.7 ± 2.9 (11.0 / 26.6) | 13.5 ± 2.8 (7.8 / 20.2) | 15.8 ± 1.4 (12.4 / 20.6) | 19.7 ± 2.9 (11.0 / 26.6) |
| | L402 | 17.3 ± 3.1 (10.8 / 24.1) | 13.6 ± 2.8 (9.9 / 20.1) | 17.0 ± 1.3 (12.5 / 21.5) | 13.7 ± 2.6 (7.1 / 22.2) |
| | P403 | 15.0 ± 3.3 (8.1 / 21.6) | 12.6 ± 2.7 (6.7 / 19.4) | 16.9 ± 1.5 (11.5 / 21.9) | 12.1 ± 3.0 (4.6 / 22.1) |
| | D404 | 13.3 ± 2.7 (7.4 / 20.1) | 9.4 ± 2.8 (3.6 / 16.4) | 13.2 ± 1.5 (8.6 / 18.0) | 9.7 ± 2.7 (3.2 / 18.9) |
| | K405 | 11.0 ± 2.1 (6.0 / 18.0) | 7.7 ± 3.1 (1.4 / 15.5) | 10.5 ± 1.6 (5.3 / 15.5) | 8.6 ± 2.6 (2.6 / 18.6) |
| | S406 | 10.3 ± 1.9 (5.6 / 20.6) | 5.7 ± 3.2 (-0.4 / 13.2) | 7.8 ± 1.5 (3.9 / 12.5) | 8.2 ± 2.4 (2.6 / 16.4) |
| | R407 | 9.6 ± 1.9 (4.5 / 17.2) | 2.1 ± 3.4 (-4.4 / 10.7) | 4.6 ± 1.5 (0.4 / 9.9) | 5.3 ± 2.4 (-0.2 / 12.9) |
| | Q408 | 10.9 ± 2.2 (5.5 / 20.6) | 1.7 ± 3.3 (-4.8 / 9.7) | 4.1 ± 1.4 (0.3 / 9.4) | 6.5 ± 2.3 (1.4 / 15.3) |
| | Q408[a] | 8.4 ± 2.6 (2.7 / 21.8) | 0.2 ± 4.2 (-8.3 / 12.0) | 2.5 ± 1.8 (-2.6 / 11.1) | 7.9 ± 2.6 (0.8 / 16.9) |
| | G409 | 14.1 ± 2.3 (8.3 / 23.0) | 2.2 ± 2.9 (-3.8 / 9.5) | 4.4 ± 1.3 (0.5 / 9.8) | 6.4 ± 2.6 (1.0 / 14.7) |
| | K410 | 16.5 ± 2.4 (11.8 / 25.3) | 5.6 ± 2.7 (-0.4 / 12.7) | 7.6 ± 1.3 (3.9 / 12.9) | 9.0 ± 2.3 (3.8 / 16.5) |
| β6-β7 loop | H449 | 23.3 ± 3.2 (15.5 / 39.0) | 3.6 ± 1.8 (-0.3 / 9.4) | 6.0 ± 1.5 (2.1 / 11.4) | 8.7 ± 4.0 (1.4 / 19.3) |
| | G450 | 24.2 ± 3.7 (15.8 / 37.2) | 2.3 ± 1.9 (-2.1 / 8.4) | 5.8 ± 1.6 (1.4 / 11.3) | 9.1 ± 4.4 (1.4 / 21.4) |
| | R451 | 22.7 ± 4.4 (13.4 / 37.8) | -1.0 ± 1.8 (-5.5 / 5.1) | 3.8 ± 2.0 (-1.1 / 11.2) | 6.6 ± 4.7 (-1.2 / 20.3) |
| | F452 | 20.8 ± 4.5 (11.3 / 34.7) | 0.2 ± 1.6 (-4.4 / 5.5) | 6.4 ± 2.5 (1.1 / 14.8) | 6.6 ± 4.1 (-1.1 / 19.4) |
| | F452[b] | 21.6 ± 5.5 (10.2 / 37.2) | -2.5 ± 1.9 (-7.7 / 4.0) | 6.1 ± 2.3 (-0.9 / 14.9) | 6.1 ± 4.1 (-2.8 / 19.9) |
| | G453 | 19.6 ± 4.7 (10.3 / 33.6) | 2.9 ± 1.4 (-1.7 / 7.2) | 8.2 ± 2.6 (2.1 / 16.6) | 7.2 ± 3.5 (0.3 / 18.6) |
| | R454 | 18.1 ± 4.4 (9.3 / 30.7) | 1.6 ± 1.5 (-1.8 / 6.3) | 5.7 ± 2.3 (0.5 / 13.4) | 5.8 ± 3.4 (-0.9 / 16.0) |

[a]Reference point taken to be Cδ.

[b]Reference point taken to be the center of mass of the Phe ring.

#### Figure S6

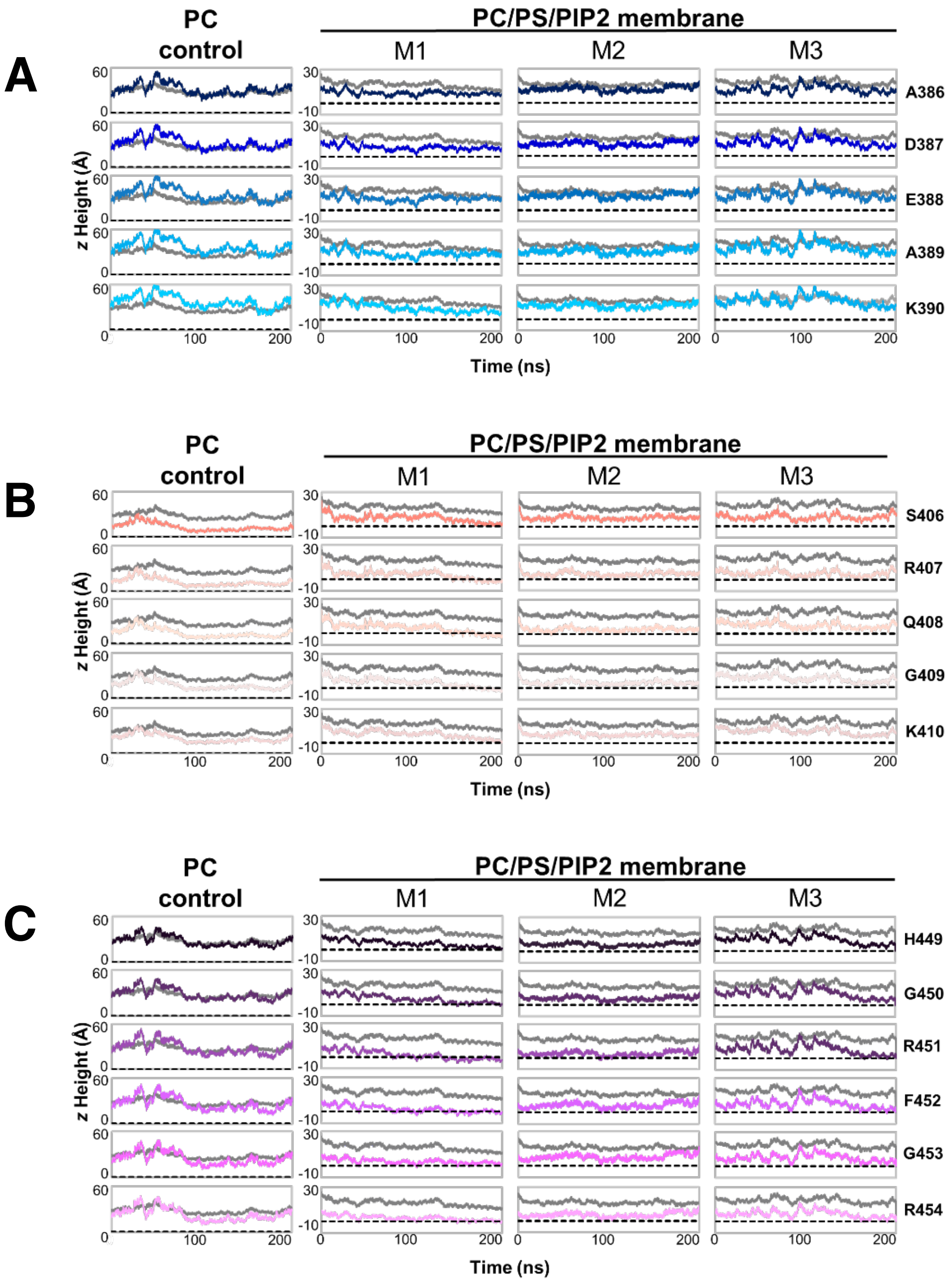

**Figure S6. Residue-level penetration calculations.** Distances of α-carbons from the membrane phosphate average position (black dashed line) are shown of individual residues over simulation time, with the protein center of mass (gray solid) for reference. Positive values are above the phosphate plane, while negative values indicate insertion. **A:** The β2-β3 loop showed no α-carbon penetration below the phosphate plane in any simulation. **B:** The β3-β4 loop had some penetration near the end of Model 1, and also had the closest contact of the three loops in the PC control. **C:** In the β6-β7 loop, the α-carbons of R451 and F452 penetrated near the membrane phosphate plane in all three PC/PS/$PIP_2$ models.

#### Figure S7

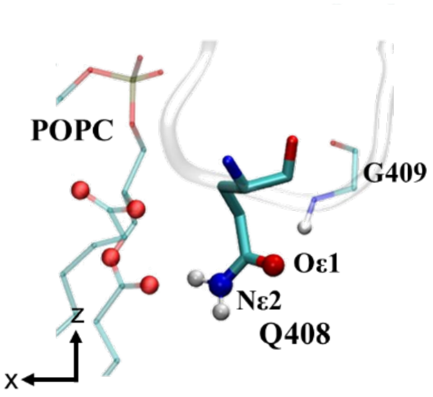

**Figure S7. Q408 sidechain H-bonding in Model 1.** The polar sidechain of Q408 (thick lines) projected toward the membrane during this simulation, and at times inserted below the phosphate plane. This insertion was stabilized through hydrogen bonding of Nε2 on Q408 (blue sphere) the ester carbonyl oxygen of a nearby POPC lipid (shown as thin lines with carbonyl oxygens as red spheres). At the same time, the arched conformation allowed Oε1 of Q408 (red sphere) to hydrogen bond with the backbone NH of G409 (shown as thin lines with the NH hydrogen as a white sphere).

#### Figure S8

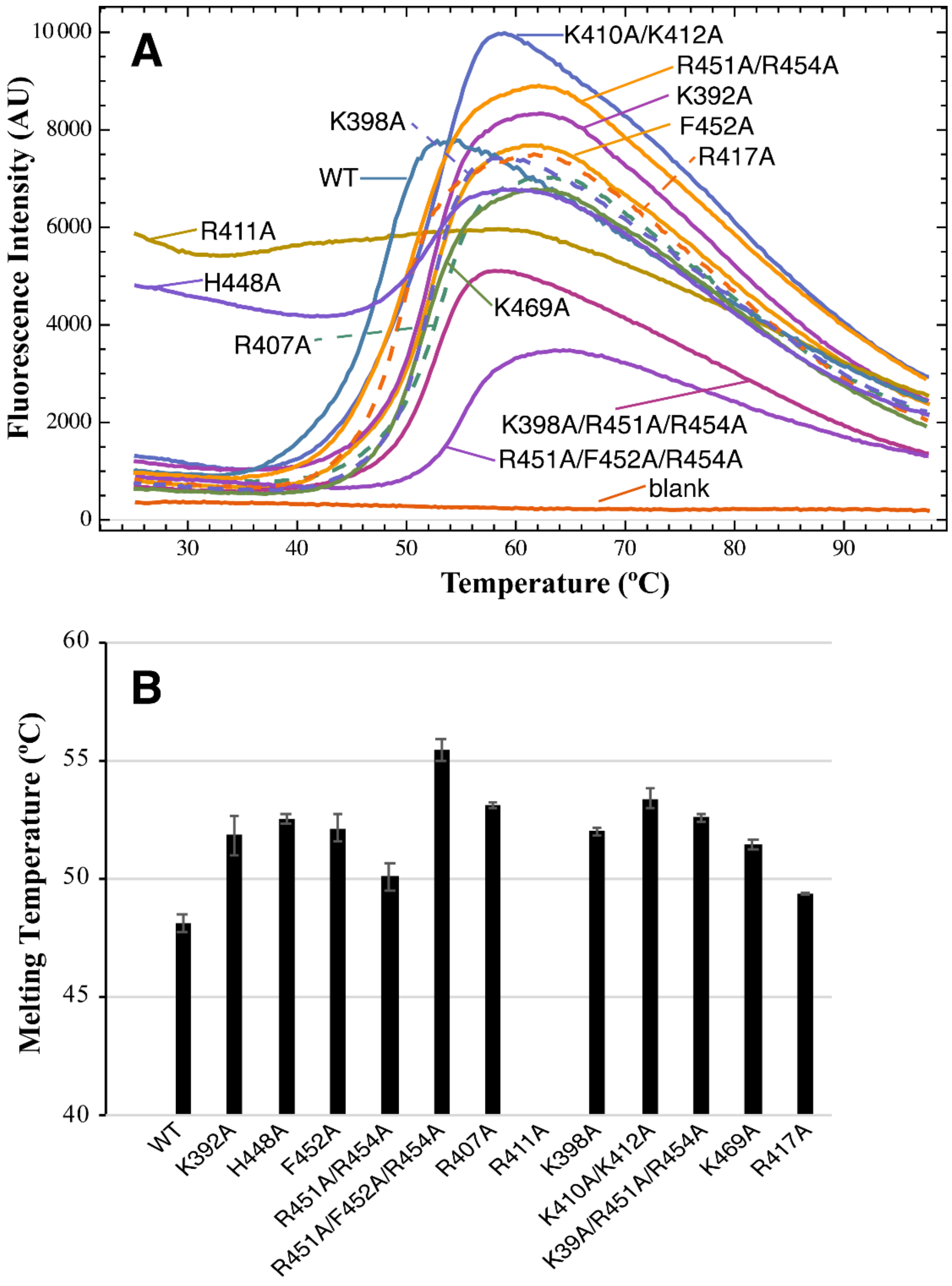

**Figure S8: Thermal stability of Slp-4 C2A mutants.** Thermal stability was measured using differential scanning fluorimetry, using a StepOne qPCR machine with the protein thermal shift dye kit from Applied Biosystems, and using 0.1 mg/ml protein. Melting temperatures were calculated as the temperature at which the slope of the melt curve was maximal. **A:** Representative melt curves for the wild-type (WT) domain and indicated mutants. **B:** Average ± S.D. from n = 4 replicate measurements. R411A did not have a clear melting transition.

**Table S5.** PS and $PIP_2$ contact for mutants compared to WT. Lipid contacts for each indicated mutant were averaged over the last 100 ns of each simulation. Two trajectories each (Sim1 and Sim2) were run for the triple mutants, differing only in a random seed. Blue coloring indicates increased contact relative to WT, while red coloring indicates decreased contact.

| | | WT | | K398A | | F452A | | K398A/R451A/R454A Sim1 | | K398A/R451A/R454A Sim2 | | R451A/F452A/R454A Sim1 | | R451A/F452A/R454A Sim2 | |
|---|---|---|---|---|---|---|---|---|---|---|---|---|---|---|---|
| | **Residue** | Avg PS Contact | Avg $PIP_2$ Contact | Avg PS Contact | Avg $PIP_2$ Contact | Avg PS Contact | Avg $PIP_2$ Contact | Avg PS Contact | Avg $PIP_2$ Contact | Avg PS Contact | Avg PIP2 Contact | Avg PS Contact | Avg $PIP_2$ Contact | Avg PS Contact | Avg PIP2 Contact |
| **β2- β3 loop** | **K390** | 0.0 ± 0.1 | 0 | 2.0 ± 0.8 | 0 | 1.5 ± 0.5 | 0 | 0.0 ± 0.0 | 0 | 1.3 ± 1.0 | 0 | 0.3 ± 0.7 | 0 | 0.0 ± 0.0 | 0 |
| | **K391** | 0 | 0 | 0 | 0 | 0.4 ± 0.5 | 0 | 0 | 0 | 0 | 0 | 0 | 0 | 0 | 0 |
| | **R392** | 0.5 ± 0.7 | 0 | 1.6 ± 0.7 | 0 | 0 | 0 | 0.0 ± 0.0 | 0 | 1.6 ± 1.1 | 0 | 0.3 ± 0.6 | 0 | 0.0 ± 0.1 | 0 |
| **β6- β7 loop** | **H449** | 0.0 ± 0.1 | 0.9 ± 0.3 | 0 | 0 | 0 | 0 | 0 | 0 | 0 | 0 | 0 | 0 | 0 | 0 |
| | **R451/A451** | 0.5 ± 0.8 | 0 | 1.5 ± 0.6 | 0 | 1.0 ± 0.0 | 0 | 0.0 ± 0.1 | 0 | 0.3 ± 0.4 | 0 | 0.4 ± 0.5 | 0 | 0.0 ± 0.2 | 0.0 ± 0.1 |
| | **R454/A454** | 0.1 ± 0.3 | 0.1 ± 0.3 | 0 | 0 | 0 | 0.0 ± 0.2 | 0.1 ± 0.3 | 0.0 ± 0.0 | 0.0 ± 0.0 | 0.1 ± 0.2 | 0.2 ± 0.4 | 0.0 ± 0.0 | 0.0 ± 0.2 | 0.3 ± 0.5 |
| **β3- β4 loop** | **K405** | 0.3 ± 0.5 | 0 | 0.1 ± 0.3 | 0 | 0.0 ± 0.1 | 0 | 1.0 ± 0.4 | 0 | 0.5 ± 0.5 | 0 | 0.2 ± 0.4 | 0.1 ± 0.2 | 0.2 ± 0.4 | 0 |
| | **R407** | 0.9 ± 0.5 | 0 | 0.8 ± 0.7 | 0 | 1.0 ± 0.3 | 0.2 ± 0.4 | 1.2 ± 0.5 | 0 | 0.7 ± 0.5 | 0 | 0.4 ± 0.7 | 0.0 ± 0.1 | 0.2 ± 0.4 | 0 |
| | **R411** | 0.7 ± 0.5 | 0 | 1.1 ± 0.4 | 0 | 0 | 0.0 ± 0.1 | 0.0 ± 0.0 | 0.1 ± 0.3 | 0.5 ± 0.6 | 0.0 ± 0.2 | 0.5 ± 0.5 | 0 | 1.5 ± 1.0 | 0 |
| **Lysine cluster** | **K398/A398** | 0.6 ± 0.5 | 1.0 ± 0.1 | 0 | 0 | 0 | 1.0 ± 0.1 | 0 | 0.0 ± 0.1 | 0 | 0.1 ± 0.3 | 0 | 1.0 ± 0.2 | 0 | 1.0 ± 0.0 |
| | **K410** | 0.4 ± 0.5 | 0 | 0.0 ± 0.2 | 0 | 0.1 ± 0.0 | 0.8 ± 0.4 | 0 | 0.9 ± 0.4 | 0.1 ± 0.3 | 0.6 ± 0.5 | 0 | 1.0 ± 0.1 | 0.1 ± 0.2 | 1.0 ± 0.2 |
| | **K412** | 0.0 ± 0.1 | 1.0 ± 0.0 | 0.1 ± 0.3 | 0 | 0 | 0.8 ± 0.4 | 0.0 ± 0.0 | 0.6 ± 0.5 | 0 | 1.0 ± 0.0 | 0.0 ± 0.0 | 0.1 ± 0.3 | 0.1 ± 0.2 | 1.0 ± 0.0 |
| **Other** | **R417** | 0.3 ± 0.5 | 0.0 ± 0.1 | 0.1 ± 0.3 | 0 | 0.2 ± 0.5 | 0 | 0.0 ± 0.1 | 0 | 0.2 ± 0.4 | 0 | 0.5 ± 0.6 | 0 | 0.0 ± 0.2 | 0 |
| | **R429** | 0.0 ± 0.0 | 0 | 0.0 ± 0.1 | 0 | 0 | 0 | 0 | 0 | 0.0 ± 0.0 | 0 | 0.2 ± 0.5 | 0 | 0.0 ± 0.0 | 0 |
| | **R440** | 0.0 ± 0.0 | 0 | 0 | 0 | 0 | 0 | 0.4 ± 0.5 | 0 | 0.1 ± 0.2 | 0 | 0.0 ± 0.1 | 0 | 0.0 ± 0.2 | 0 |
| | **K483** | 0.0 ± 0.0 | 0 | 0.0 ± 0.1 | 0.5 ± 0.5 | 0.0 ± 0.0 | 0 | 0.0 ± 0.0 | 0.0 ± 0.0 | 0.0 ± 0.2 | 0 | 0 | 0 | 0 | 0 |
| | **Total** | **4.2 ± 1.6** | **3.0 ± 0.5** | **7.4 ± 1.6** | **0.5 ± 0.5** | **4.3 ± 0.9** | **2.8 ± 0.7** | **2.7 ± 0.9** | **1.5 ± 0.7** | **5.1 ± 1.9** | **1.7 ± 0.6** | **2.9 ± 1.7** | **2.1 ± 0.5** | **2.2 ± 1.3** | **3.3 ± 0.5** |

= significant (at least 1 σ) increase in contact*
= significant (at least 1 σ) decrease in contact*
= moderate (0.5 – 1 σ) increase in contact*
= moderate (0.5 – 1 σ) decrease in contact*

*All mutants compared to WT

### Figure S9

**kTSIK (k=phosphogluconoyl-lysine) MSMS fragment spectrum, MH+ = 834.3850**

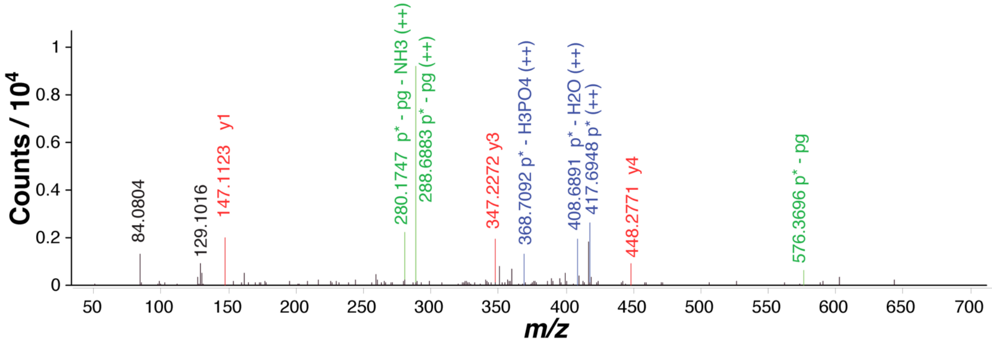

**Peak List**

| Obs. m/z | Pred. m/z | ID |
|---|---|---|
| 84.0804 | 84.0808 | lysine fragment |
| 129.1016 | 129.1022 | lysine |
| 147.1123 | 147.1128 | y1 |
| 280.1747 | 280.1761 | p* - pg - NH3 (++) |
| 288.6883 | 288.6894 | p* - pg (++) |
| 347.2272 | 347.2289 | y3 |
| 368.7092 | 368.7085 | p* - H3PO4 (++) |
| 408.6891 | 408.6911 | p*-H2O (++) |
| 417.6948 | 417.6964 | p* (++) |
| 448.2771 | 448.2766 | y4 |
| 576.3696 | 576.3715 | p* - pg |

**KTSIK (unmodified) MSMS fragment spectrum, MH+ = 576.3721**

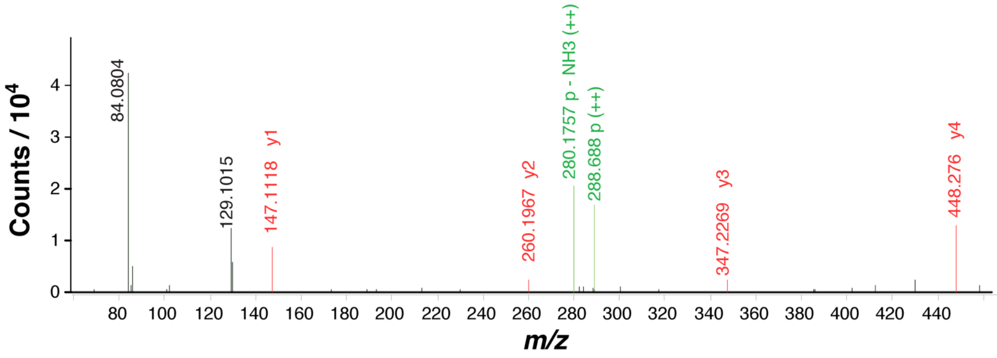

**Peak List**

| Obs. m/z | Pred. m/z | ID |
|---|---|---|
| 84.0804 | 84.0808 | lysine fragment |
| 129.1015 | 129.1022 | lysine |
| 147.1118 | 147.1128 | y1 |
| 260.1967 | 260.1969 | y2 |
| 280.1757 | 280.1761 | p - NH3 (++) |
| 288.688 | 288.6894 | p (++) |
| 347.2269 | 347.2289 | y3 |
| 448.276 | 448.2766 | y4 |

**Figure S9: Identification of probable phosphogluconoylation at K412.** ESI-MS-MS was performed on a tryptic digest of the early peak, and peptides corresponding to modified and unmodified tryptic fragments were identified via molecular formula matching. Shown are MSMS spectra of peptides with the indicated masses (MH+), corresponding to kTSIK$^{412\text{-}416}$ (top), where k is phosphogluconoyl-lysine, and KTSIK$^{412\text{-}416}$ (bottom). Peaks present in the kTSIK but not KTSIK spectrum correspond to the kTSIK precursor ion (p*) and losses of water and water+phosphate (blue). (Note, the loss of phosphate peak confirms that the phosphate is distal to the gluconoyl group.) Although b-ion fragments are not visible, the y-ion series shows the same pattern in both spectra (red), indicating that the modification is on K412. The KTSIK precursor ion mass (p) is also visible in the kTSIK spectrum consistent with loss of phosphogluconate (p* - pg) (green). ++ indicates $z = 2$.
